# Horizontally acquired quorum sensing regulators recruited by the PhoP regulatory network expand host-adaptation repertoire in the phytopathogen *Pectobacterium carotovorum*

**DOI:** 10.1101/776476

**Authors:** Daniel Bellieny-Rabelo, Ntombikayise Precious Nkomo, Divine Yufetar Shyntum, Lucy Novungayo Moleleki

## Abstract

In this study, we examine the impact of transcriptional network rearrangements driven by horizontal gene acquisition in PhoP and SlyA regulons using as a case study the phytopathosystem comprised of potato tubers and the soft rot pathogen *Pectobacterium carotovorum* subsp. *brasiliense* (*Pcb*1692). By comparing those two networks with that of PecS obtained from the closely related *Dickeya dadantii*, we found that: (a) 24-31% of the genes regulated at late infection are genus-specific (GS) (based on Pectobacterium and Dickeya genera), and that (b) of these, 28.1-44.4% were predicted with high confidence as horizontal gene transfer (HGT) candidates. Further, genome simulation and statistical analyses corroborated the bias in late infection regulons towards the transcriptional control of candidate GS-HGT genes by PhoP, SlyA, and PecS, highlighting the prominence of network rearrangements in these late infection regulons. The evidence further supports the circumscription of two horizontally acquired quorum sensing regulators (*car*R and *exp*R1) by the PhoP network. By recruiting *car*R and *exp*R1, the PhoP network also impacts certain host adaptation- and bacterial competition-related systems, seemingly in a quorum sensing-dependent manner, such as the type VI secretion system, carbapenem biosynthesis, and plant cell walls degrading enzymes (PCWDE) such as cellulases and pectate lyases. Conversely, polygalacturonases and the type III secretion system (T3SS) exhibit a transcriptional pattern that suggests quorum sensing-independent regulation by the PhoP network. This includes a yet uncharacterized novel phage-related gene family within the T3SS gene cluster that has been recently acquired by two Pectobacterium species. The evidence further suggests a PhoP-dependent regulation of carbapenem and PCWDE-encoding genes based on the synthesized products’ optimum pH. The PhoP network also controls *sly*A expression *in planta*, which seems to impact the carbohydrate metabolism regulation, especially at early infection when 69.6% of the SlyA-regulated genes from that category also require PhoP to achieve normal expression levels.

**AUTHOR SUMMARY:** Exchanging genetic material through horizontal transfer is a critical mechanism that drives bacteria to efficiently adapt to host defenses. In this report, we demonstrate that a specific plant pathogenic species (from the Pectobacterium genus) successfully integrated a population density-based behaviour system (quorum sensing) acquired through horizontal transfer into a resident stress-response gene regulatory network controlled by the PhoP protein. Evidence found here underscores that subsets of bacterial weaponry critical for colonization, typically known to respond to quorum sensing, are also controlled by PhoP. Some of these traits include different types of enzymes that can efficiently break plant cell walls depending on the environmental acidity level. Thus, we hypothesize that PhoP ability to elicit regulatory responses based on acidity and nutrient availability fluctuations may have strongly impacted the fixation of its regulatory connection with quorum sensing. In addition, another global gene regulator known as SlyA was found under the PhoP regulatory network. The SlyA regulator controls a series of carbohydrate metabolism-related traits, which also seem to be regulated by PhoP. By centralizing quorum sensing and *sly*A under PhoP scrutiny, Pectobacterium cells added an advantageous layer of control over those two networks that potentially enhances colonization efficiency.

## INTRODUCTION

Highly specialized colonization traits are constantly selected in phytopathogenic gram-negative bacteria in order to overcome severe obstacles imposed by plant apoplasts. The apoplastic space comprises a nutrient-poor milieu that harbors an extensive inventory of toxic and defense-related molecules, such as plant antimicrobial peptides, reactive oxygen species, and plant organic compounds (1, 2). In addition, the apoplastic space is generally acidic (ranging between pH 4.5-6.5) due to the presence of organic acids such as malic and citric acids, which also enforces the anti-growth strategy against pathogenic invasions (3). In this context, a crucial regulatory system in bacteria, which is frequently associated with response to acidic stress, is the PhoQ/PhoP two-component system (4). This response-based system is highly conserved and widely studied across bacterial lineages because of its prominent role as a global transcriptional regulator.

The simultaneous flow of transcription/translation in bacterial cells inhabiting such competitive environments requires highly optimized regulatory mechanisms to ensure accurate responses during colonization. Thus, in order to cope with frequent gene gains and losses that result mainly from horizontal transfer events, bacterial regulatory circuits must efficiently accommodate newly acquired genes (5). Horizontal gene transfer is perceived as a critical driving force in the evolution of bacterial genomes, from which a number of pathogenicity themes emerge, including some secretion systems and prophages (6, 7). Hence, it is not surprising that the influence of important virulence regulators such as SlyA (8) or PecS (9) over regions incorporated through HGT have been previously reported. Both SlyA and PecS belong to the MarR family of transcriptional regulators, which encompasses a range of genes involved in virulence and antibiotic resistance control (10–12).

Among gram-negative bacteria, one specific group commonly referred to as soft rot Pectobacteriaceae (SRP) (13, 14) (formerly known as soft rot Enterobacteriaceae), has increasingly gained attention over the last few decades as causative agents of wilt/blackleg diseases in economically important crops worldwide (15, 16). This group is most prominently represented by Pectobacterium and Dickeya genera. The SRPs are opportunistic gram-negative pathogens capable of producing distinctively high amounts of pectinolytic enzymes compared to other pectolytic bacteria (e.g. *Clostridium* spp., *Bacillus* spp., *Pseudomonas* spp.) (17). These plant cell wall degrading enzymes encompass a variety of families that concertedly promote disease through tissue maceration (18). While some PCWDE classes exhibit close to neutral or high optimum pH, such as cellulases, pectate and pectin lyases (Cel; Pel; Pnl), others function at low optimum pH, namely polygalacturonases (Peh) (19, 20). In this sense, the expression of different groups of PCWDEs tends to be regulated according to the pH within the plant tissue, which is acidic in the apoplast at first, then becomes progressively basic as the disease progresses (17). In this context, one of the best-characterized mechanism of PCWDE regulation in SRP pathogens is the quorum-sensing (QS) (21). This is a crucial regulatory circuit controlling population density-based behaviour in SRP, as well as in a large spectrum of bacteria (22, 23). Quorum sensing networks in gram-negative bacteria rely on recognition of autoinducer molecules, such as acyl-homoserine lactones (AHL) and other products synthesized using S-adenosylmethionine, by specialized receptors such as ExpR/LuxR homologs (24, 25). The complexed form, ExpR/LuxR-autoinducer, is then able to bind specific DNA regions in order to participate in transcriptional regulation (26). One of the regions regulated by the ExpR-autoinducer complex is the *rsm*A promoter region. Thereafter, the transcriptional repression of *rsm*A prevents RsmA-induced inactivation of PCWDE (27). In addition to PCWDEs, QS has been shown to regulate other traits involved in virulence and interbacterial competition, such as the carbapenem antibiotic (*car* genes) and the type III and VI secretion systems (T6SS) in Pectobacterium spp. (21, 28).

In the present report, we investigate the impact of transcriptional network rearrangements in two global transcriptional regulators (PhoP and SlyA) over the *in planta* regulation of crucial host adaptation- and fitness-oriented systems. Since regulatory network rearrangement events have the potential to give rise to phenotypic differences in pathogenesis development among these organisms, understanding these processes is crucial to unveiling how these networks shape pathogenicity in different lineages.

## RESULTS

### RNA-Seq mapping and the reach of PhoP and SlyA regulons *in planta*

In order to examine both PhoP and SlyA regulatory networks *in planta*, we engineered *Pcb*1692Δ*sly*A and *Pcb*1692Δ*pho*P mutants using the lambda recombination technique (see ‘Methods’ for details). The integrity of the mutants and complement strains were confirmed by PCR analyses (S1 Fig and S2 Fig) as well as by DNA sequencing. Additionally, *in vitro* growth assay demonstrated that the deletion of either *Pcb*1692Δ*pho*P or *Pcb*1692Δ*sly*A genes did not impair the growth of the mutant strains (S3 Fig and S4 Fig). Next, the global impact caused by *pho*P or *sly*A deletion in the *Pcb*1692 genome towards *in planta* transcriptional profiles were analyzed in original whole-transcriptome data sets comprising ∼124 million RNA-Seq reads (Table 1). By comparing significant gene expression changes between wild-type and either *Pcb*1692Δ*sly*A or *Pcb*1692Δ*pho*P mutants, we infer transcriptional regulation affected by these regulatory networks (see ‘Methods’ for details). Additional validation by qRT-PCR of the differentially expressed genes of particular interest for this study was also conducted (S5 Fig). The samples selected for this study depicted two stages of plant infection: 12- and 24-hours post-infection (hpi) denoted here as early (EI) and late infection (LI), respectively. The dataset exhibited good quality, as 89.2% (∼111 million) of the reads were mapped on the *Pcb*1692 reference genome (Table 1). Of these, 88.9% (∼98 million reads) were uniquely mapped on the reference genome. Transcriptional variations between wild-type *Pcb*1692 and mutant strains during infection enabled collective identification of 511 genes affected (up-/down-regulation; see ‘Methods’ for details) by both PhoP and SlyA regulatory networks in the two time points (S1 Table).

**Table 1:**
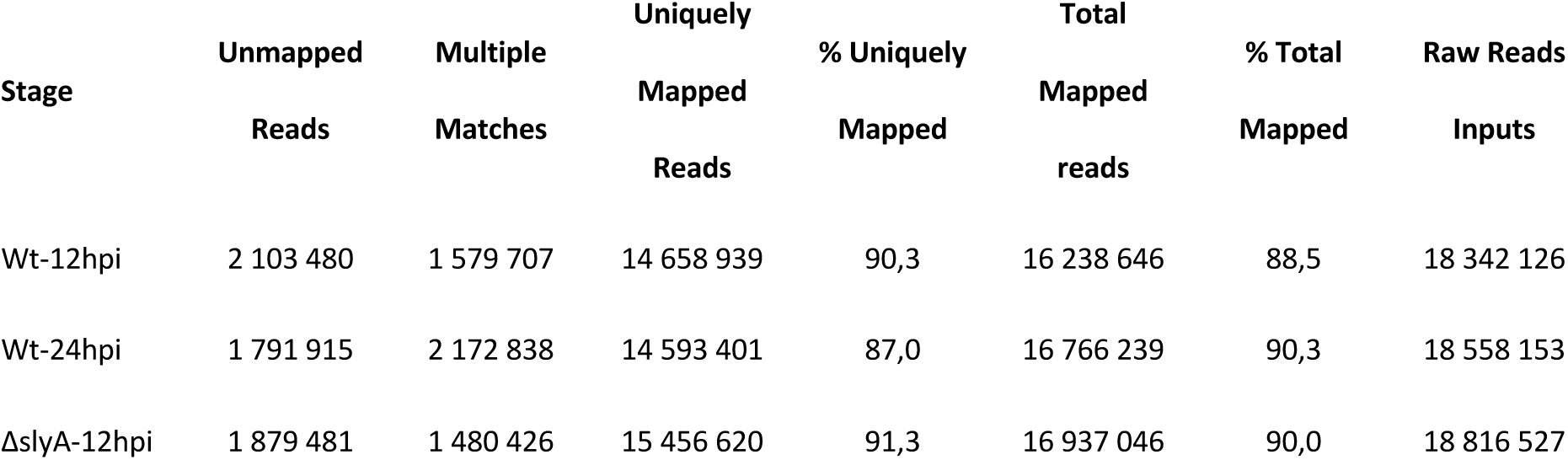
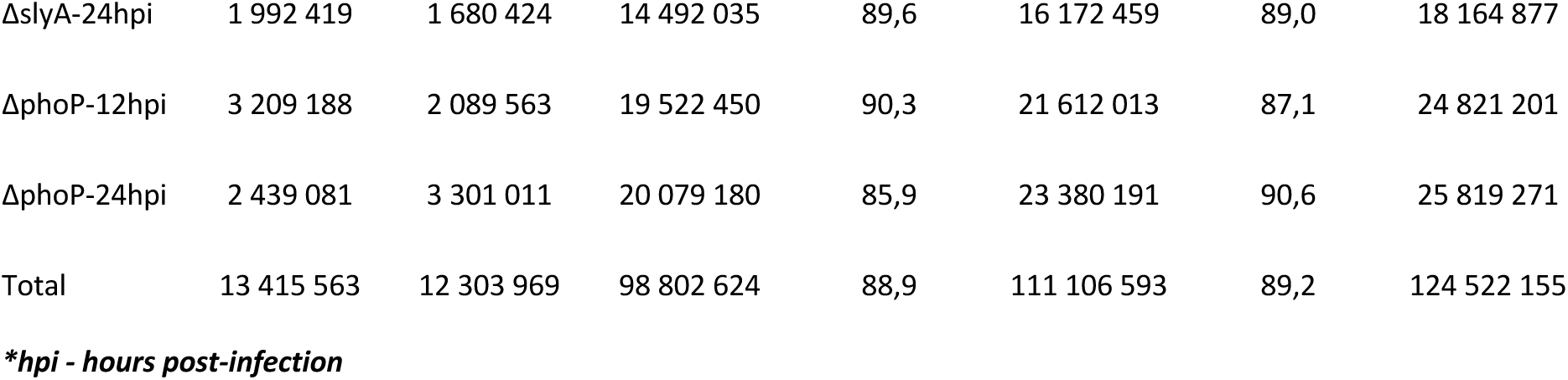
Mapping summary of RNA-Seq reads from mutant strains on *Pcb*1692 reference genome.

| Stage | Unmapped<br>Reads | Multiple<br>Matches | Uniquely<br>Mapped<br>Reads | % Uniquely<br>Mapped | Total<br>Mapped<br>reads | % Total<br>Mapped | Raw Reads<br>Inputs |
| --- | --- | --- | --- | --- | --- | --- | --- |
| Wt-12hpi | 2 103 480 | 1 579 707 | 14 658 939 | 90,3 | 16 238 646 | 88,5 | 18 342 126 |
| Wt-24hpi | 1 791 915 | 2 172 838 | 14 593 401 | 87,0 | 16 766 239 | 90,3 | 18 558 153 |
| ΔslyA-12hpi | 1 879 481 | 1 480 426 | 15 456 620 | 91,3 | 16 937 046 | 90,0 | 18 816 527 |
| ΔslyA-24hpi | 1 992 419 | 1 680 424 | 14 492 035 | 89,6 | 16 172 459 | 89,0 | 18 164 877 |
| ΔphoP-12hpi | 3 209 188 | 2 089 563 | 19 522 450 | 90,3 | 21 612 013 | 87,1 | 24 821 201 |
| ΔphoP-24hpi | 2 439 081 | 3 301 011 | 20 079 180 | 85,9 | 23 380 191 | 90,6 | 25 819 271 |
| Total | 13 415 563 | 12 303 969 | 98 802 624 | 88,9 | 111 106 593 | 89,2 | 124 522 155 |

**\*hpi - hours post-infection**

From the PhoP regulon, we detected a 30.7% reduction in the total network size between early and late infection (Figure 1A). This phenomenon could result from a variety of non-mutually exclusive events, among which the most predictable would be: (a) increased number of active transcriptional regulators at 24 hpi competing with PhoP for promoter regions binding, (b) PhoP dephosphorylation by PhoQ following the absence of triggering conditions, or (c) decreased transcription of the *pho*P gene. Thus, to further inspect this shift in PhoP network sizes between EI and LI, we set out to evaluate the transcriptional variation of the *pho*P gene *in planta* by *Pcb*1692 wild type. First, *Pcb*1692 was inoculated into potato tubers and samples were harvested at 8, 12, 16, 20 and 24 hpi. Next, qRT-PCR was used to analyze the expression of the *pho*P gene in those samples. The results showed that the relative expression of *pho*P gene nearly doubled between 8 and 12 hpi (Figure 1B). This increase was followed by a significant decrease (p-value = 0.042) between 12 and 16 hpi. After keeping a constant expression level between 16 and 20 hpi, *Pcb*1692 cells drastically decrease *pho*P gene expression at 24 hpi (p-value = 0.004). This converges with the shrinkage observed in the network size. Hence, since PhoP is frequently associated with transcriptional responses to acidic environments and low Mg^2+^ concentration, the elevated *pho*P transcription in the early stages of infection is arguably a direct response to the nutrient poor environment found in the apoplast. Conversely, after a certain time point between 20 and 24 hpi, *pho*P transcription decreases, which is consistent with the reduced cellular demand for this regulator following environmental alkalization due to progressive host cell lysis.

**Figure 1.**
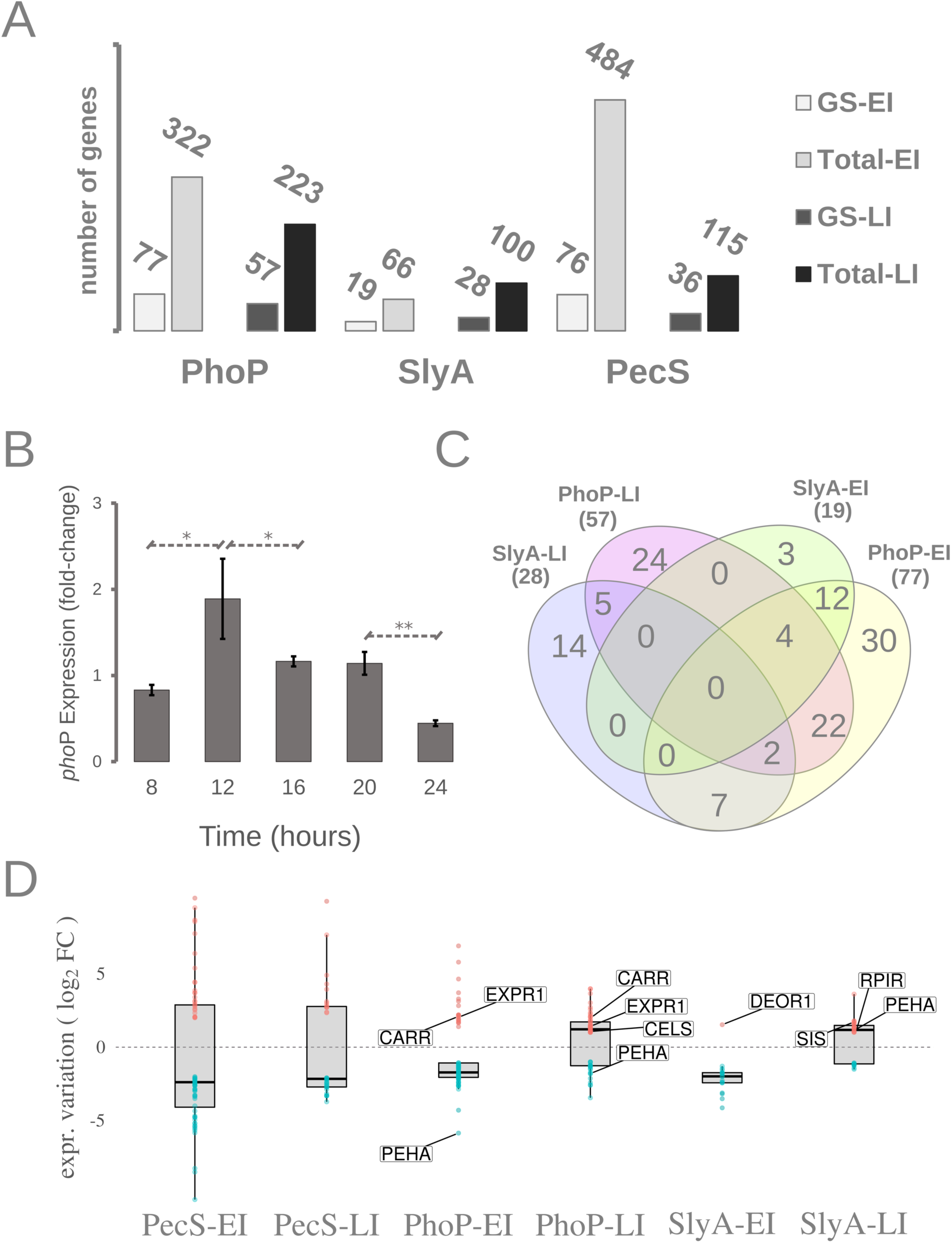
Genome-wide regulatory impact of three global regulators in *Pcb*1692 and *Ddad*3937 and the transcriptional levels of *pho*P regulator during plant infection. **(A)** Genus-specific (GS) versus the total gene (Total) amounts in PhoP, SlyA and PecS regulons *in planta* both at early and late infection (EI; LI) are represented in the bar plot. PhoP and SlyA regulons were extracted from *Pcb*1692 infection samples. PecS regulons were extracted from *Ddad*3937 samples. **(B)** Transcriptional levels of the *pho*P gene measured by qRT-PCR in four-hour intervals in potato tubers infected by *Pcb*1692 across the first 24 hpi. Consecutive time points were tested for statistically significant (t-test) differences in their expression levels. Significant results are marked with asterisks in the graph (* for *P* < 0.05; ** for *P* < 0.005). **(C)** Venn diagrams depicting genus-specific genes found in PhoP and SlyA *in planta* regulons at EI and LI assessed in *Pcb*1692. The total number of genus-specific genes found within each regulon is highlighted under each regulon label. **(D)** Gene transcriptional variations (log2 fold change) are depicted specifically for genus-specific genes. Transcriptional variation data was extracted from PhoP, SlyA and PecS regulons either at early or late infection (EI; LI). The box plots are overlaid with dot plots, in which each dot represents a single gene expression value. In PhoP and SlyA regulons, specific genes are labelled as follows: polygalacturonase A (PEHA – *PCBA_RS10070*), cellulase S (CELS – *PCBA_RS08365*), quorum sensing regulators (CARR – *PCBA_RS04390*; EXPR1 – *PCBA_RS15665*), transcriptional regulators (DEOR1 – *PCBA_RS02575*; RPIR – *PCBA_RS22175*), and SIS-encoding gene (*PCBA_RS22170*).

### Determining the genus-specific content in PhoP, SlyA, and PecS *in planta* regulons

Next, the prevalence of distinctive traits (genus-specific) was inspected within the full extent of transcriptionally regulated genes by three global regulators frequently associated with virulence and/or stress responses, namely: PhoP and SlyA in *Pcb*1692, and PecS in *Dickeya dadantii* strain 3937 (*Ddad*3937). To achieve that, we analyzed 6 distinct regulons obtained from those 3 global regulators in two different stages (i.e. early and late) of plant infection. Hence, this analysis focus on three early infection regulons, i.e. PhoP-, SlyA- and PecS-EI, along with three late infection regulons, i.e. PhoP-, SlyA-, and PecS-LI. This includes (a) a publicly available whole-transcriptome data set which utilized *Ddad*3937Δ*pec*S mutant strain (9), along with (b) two original data sets featuring *Pcb*1692Δ*sly*A and *Pcb*1692Δ*pho*P strains (NCBI Accession: PRJNA565562). The identification of distinctive systems between the two genera was achieved by predicting genus-specific genes in Pectobacterium and Dickeya genera based on protein sequence orthology. This analysis was underpinned by an extensive correlational database previously generated including 39 and 61 strains from Dickeya and Pectobacterium genera respectively (6). By using this database, we inquired for each gene in a given strain, how many orthologs could be detected in other strains within the genus, as well as in the opposing genus. Those gene products for which no orthologous counterparts exist in the opposing genus were considered, given the scope of this study, as GS in the SRP context, i.e. Pectobacterium-exclusive or Dickeya-exclusive.

Next, the presence of GS protein-coding genes was surveyed within each of the *in planta* regulons. The preliminary aim was to measure the relative presence of GS genes in these regulons in specific infection stages. Our screening revealed that across the three early infection regulons (from PhoP, SlyA, and PecS), 15-29% of the regulated genes are GS. Interestingly, at late infection, two out of three regulons (except for SlyA) exhibited increased relative GS content, ranging between 25-31% (Figure 1A). Notably, the PecS regulons exhibited the most drastic shift in the GS content proportion between EI (15%) and LI (31%). Furthermore, by comparing the GS gene sets of SlyA and PhoP regulons from *Pcb*1692, the presence of GS genes that are exclusive to individual regulons (regulon-exclusive) was more conspicuous in those of late infection. Thus, in PhoP and SlyA data sets, 44-50% (24/57; 14/28) of the GS genes regulated at LI are regulon-exclusive (Figure 1C). Conversely, 16-40% of the GS (3/19; 30/77) were regulon-exclusive among EI samples. This analysis revealed that at late infection, PhoP, SlyA (in *Pcb*1692) and PecS (in *Ddad*3937) tend to mobilize transcription of an equal or greater proportion of GS genes compared to early infection. Additionally, it also showed that PhoP and SlyA exhibit bigger relative amount of GS regulon-exclusive genes at late infection in comparison to early infection.

In the SlyA-EI data set, the only GS gene displaying increased expression was a MarR transcriptional regulator homolog to *deo*R (*PCBA_RS02575*), which is frequently associated with repression of carbohydrate metabolism (Figure 1D). Additionally, a *rpi*R homolog, which comprises another important player in the regulation of carbohydrate metabolism was also found under SlyA network exclusively at late infection, further corroborating SlyA role as a relevant player in cellular carbohydrate regulation. Interestingly, both *deo*R and *rpi*R homologs in *Pcb*1692 are regulon-exclusive respectively from SlyA-EI and SlyA-LI. This indicates a particular demand for the transcriptional regulation of certain carbohydrates metabolism-related systems controlled by these regulators in specific stages of infection.

Moreover, two Pectobacterium-exclusive genes respectively encoding a polygalacturonase (Peh) and an endoglucanase/cellulase (Cel) were found within PhoP *in planta* regulons (Figure 1D). Converging with the evidence for the transcriptional regulation of those PCWDE encoding genes, especially by PhoP, we also found two QS regulators under PhoP control, namely *exp*R1 and *car*R (Figure 1D). These QS regulators are GS genes in Pectobacterium genomes. While ExpR1 is often, but not exclusively, associated with transcriptional regulation of pectinolytic enzymes (21, 29), CarR is associated with regulation of the carbapenem antibiotic biosynthesis (28). The presence of *lux*R/*exp*R homologs within the PhoP network might be directly linked to the presence of two bacterial secretion systems. This type of association may directly impact fitness which will be inspected in a subsequent section.

### Horizontal acquisitions in the genus-specific contents and their regulation by PhoP, SlyA and PecS

While the presence of GS genes across the six regulons was observed, their origin was still unclear. The GS genes observed here may have emerged through different processes such as (a) gene loss in the neighboring lineage, (b) intra-lineage functional divergence following gene duplication, or (c) gene acquisition via HGT. Hence, aiming to evaluate the prevalence of transcriptional network rearrangement processes, especially in Pcb1692 networks, we set out to predict HGT candidates using parametric methods. As per previous observations (30), we chose a gene-based method (GC content at third-codon position - GC3) and a window-based method (dinucleotide frequencies - DINT_KL) to evaluate sequence composition bias in *Pcb*1692 and *Ddad*3937 genes (see ‘Methods’ for details). These predictions will be further compared to the orthology-based prediction of genus-specific genes as an additional layer of evidence.

By examining the entire set of GS genes in *Pcb*1692 (846 genes) we found that respectively 43.8 and 22.7% were predicted as HGT candidates by GC3 or DINT_KL methods. Importantly, 55.2% of the high-confidence HGT candidates, meaning those supported by both GC3 and DINT_KL, in *Pcb*1692 belong to the GS gene set. On the other hand, in *Ddad*3937 37.8% of the high-confidence HGT candidates are GS genes. This preliminary inquiry revealed that the GS content in *Pcb*1692 seems to include a larger portion of horizontally acquired genes compared to *Ddad*3937. Furthermore, the inspection of PecS *in planta* regulons at early and late infection in *Ddad*3937 revealed that respectively 6.6% and 44.4% of the GS genes are high-confidence HGT candidates. On the other hand, by examining PhoP and SlyA *in planta* regulons in *Pcb*1692, the high-confidence candidates within the GS genes accounted respectively for 24.7% and 10.5% at early infection, and 28.1% and 28.6% at late infection. In this context, the genus-specific HGT candidate (GS-HGT) genes most likely arise from recent gene acquisitions in a given genus. Hence, this preliminary analysis supports that the regulation of some distinctive traits observed among Pectobacterium and Dickeya genera involves recent rearrangements in PhoP, SlyA and PecS regulatory networks driven by HGT.

Thereafter, in order to inspect whether these regulatory networks are biased towards or against controlling GS-HGT candidate genes, we conducted a statistical enrichment analysis. The results from Fisher’s exact tests showed that, for the three LI regulons analyzed (from PhoP, SlyA, and PecS), there is an overrepresentation of GS-HGT genes (Figure 2A). Curiously, the PecS-EI regulon exhibits a negative correlation with GS-HGT genes. Next, we aimed to further confirm these results using a different approach. Computational simulations were performed to shuffle the genomes from each organism (i.e. *Pcb*1692 and *Ddad*3937) generating ‘pseudo-genomes’. This strategy aims to empirically evaluate the frequency to which GS-HGT genes should be expected by chance in these regulons and compare this with the real data. For each of the six regulons analyzed, this experiment generated 10,000 shuffled pseudo-genomes (see ‘Methods’ for details). The results corroborated, by empirical means, what statistical analysis had previously established (Figure 2B). Overall, these analyses showed that transient shifts, in this case, due to temporal variation, in regulon profiles can alter their bias towards the presence of recently acquired genes. Although at late infection all regulons exhibited a consistent bias towards the regulation of GS-HGT genes, none of them displayed similar bias at early infection. Contrarily, the PecS network seemed to avoid transcriptional interactions with GS-HGT genes at early infection. Overall, this evidence suggests a particular requirement for expression of distinctive traits seemingly acquired through HGT in these organisms at the late stages of infection.

**Figure 2.**
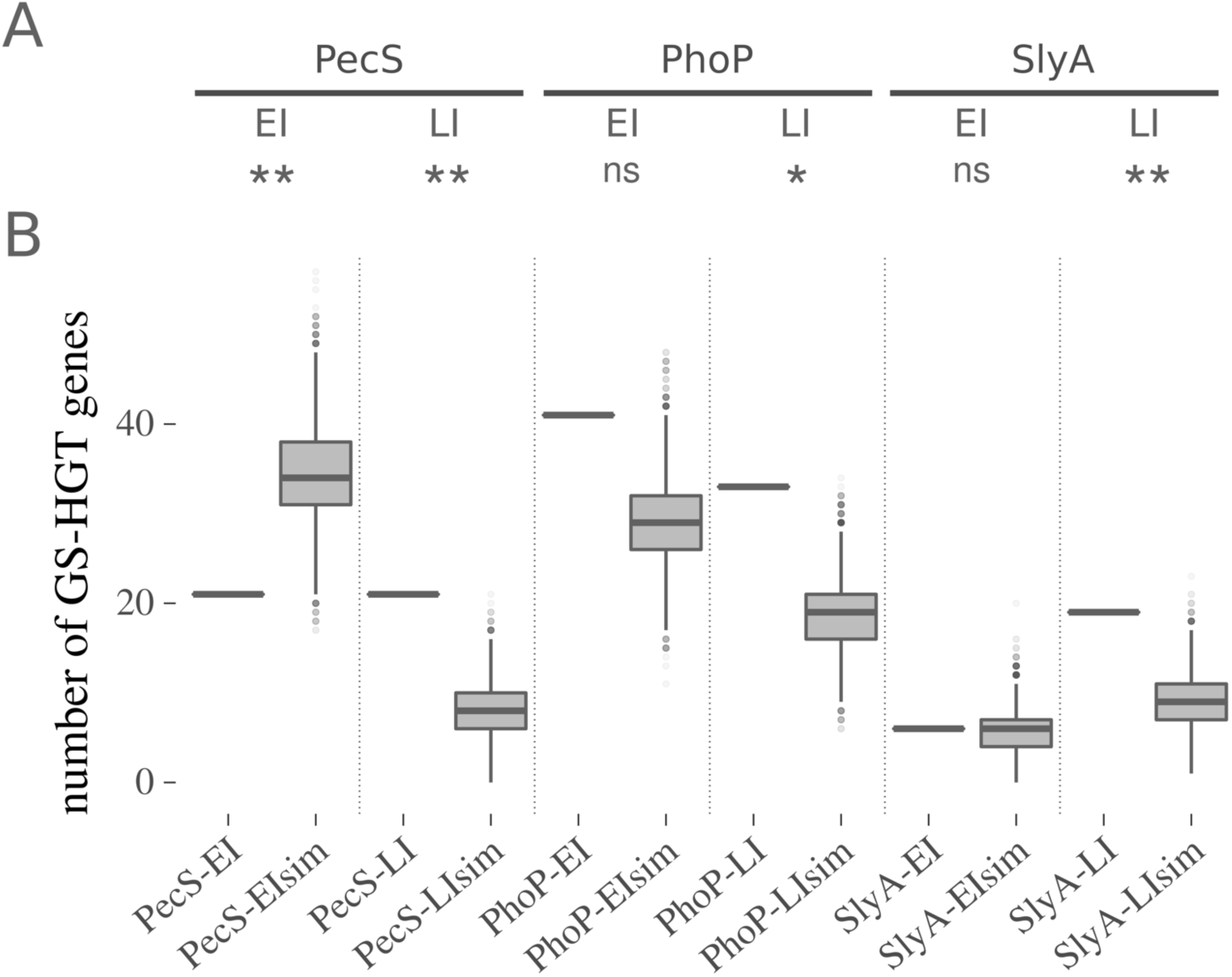
Overrepresentation of genus-specific HGT candidate genes in PhoP, SlyA and PecS regulons in *Pcb*1692 and *Ddad*3937. **(A)**The significance of genus-specific HGT candidate (GS-HGT) genes occurrence in each regulon was analyzed through two-tailed Fisher exact tests using the respective *spp*. genomes as background set (* for *P* < 0.05; ** for *P* < 0.01; ns for *P >* 0.05). **(B)** Each section of the plot (separated by dotted lines) represents the comparison between (a) the actual number of GS-HGT genes found in each regulon as a single horizontal line, and (b) the distribution of GS-HGT gene amounts found in each of the 10 000 computationally shuffled versions of the respective regulons (see ‘Methods’ for details). The simulated data sets are labelled with the ‘Sim’ postfix in the graph.

### Host adaptation and regulatory systems acquired by PhoP and SlyA networks

Highly conserved genes among the SRPs such as *pho*P (PCBA_RS01290*; DDA3937_RS11500*) or *pec*S (PCBA_RS02985*; DDA3937_RS20885)* exhibited similar overall sequence composition to their resident genomes (*Ddad*3937 or *Pcb*1692) (Figure 3A and B; S2 Table). Unlike these two regulators, *sly*A (*PCBA_RS02460; DDA3937_RS12595*) homologs consistently exhibited deviant GC3 indexes compared to the two resident genomes analyzed, although in terms of dinucleotide frequencies a significant distance was not observed. Thus, considering the conservation of *sly*A across all SRP genomes, it is likely that this gene acquisition occurred at least in the last common ancestor between Pectobacterium and Dickeya genera. Next, we observed that the SlyA-regulated GS transcriptional regulator *deo*R1 (PCBA_RS02575) had no support from HGT prediction. This indicates that this gene was likely lost in the Dickeya lineage, whereas most of the Pectobacterium retained it. In contrast, two other SlyA-regulated genes involved in the transcriptional regulation of carbohydrate metabolism, i.e. *rpi*R (*PCBA_RS22175*) and an SIS domain-encoding (*PCBA_RS22170*), were predicted as HGT candidates with high confidence. This may indicate that the Pectobacterium genus benefits from increasing the regulatory complexity of carbohydrate metabolism either through gene gain or by simply keeping ancestral transcriptional regulators in the lineage. This notion is further corroborated by the enrichment of genes associated with carbohydrate metabolism in the SlyA-EI regulon (S6 Figure).

**Figure 3.**
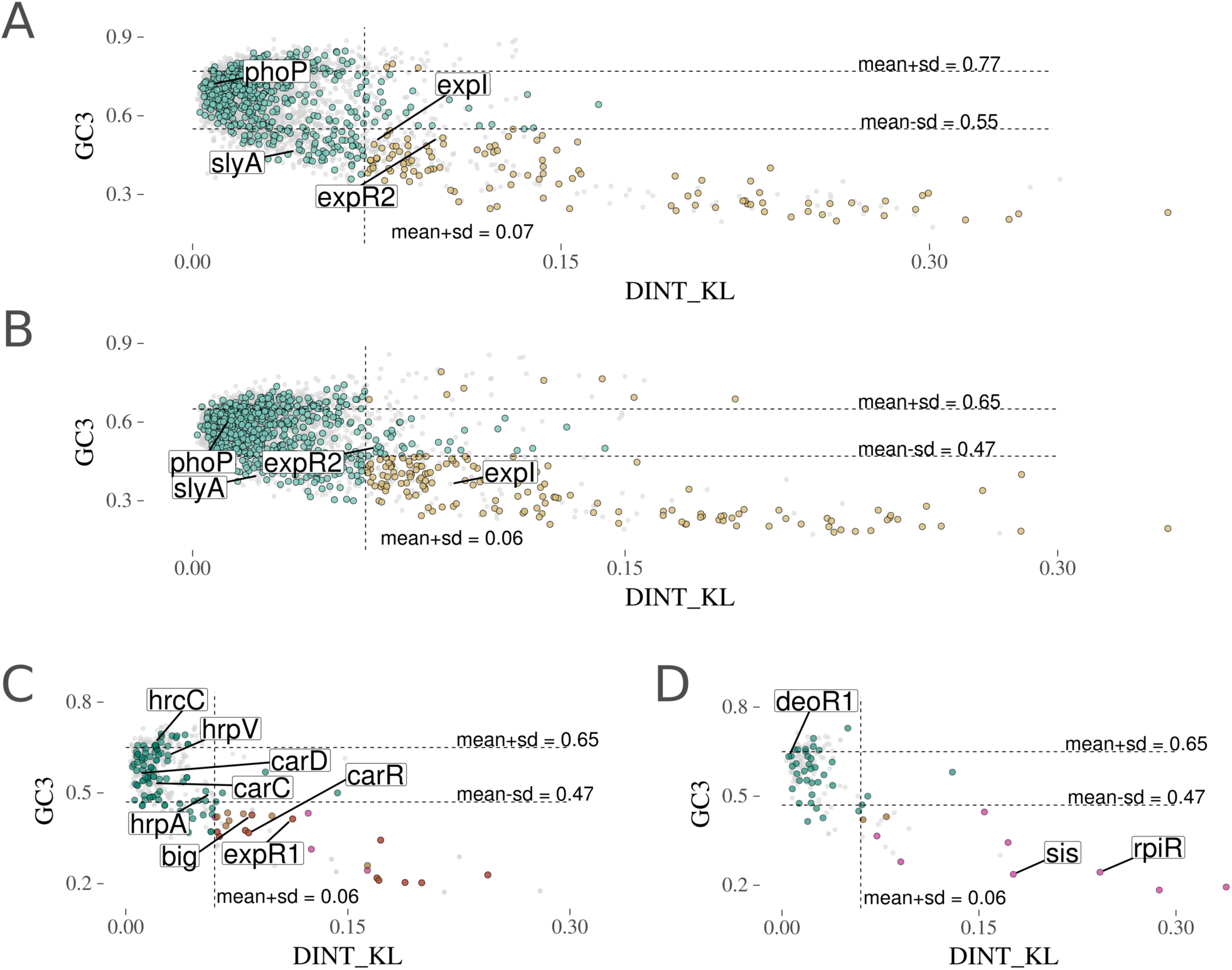
Horizontal gene transfer predictions for genus-specific genes in *Ddad*3937 and *Pcb*1692. **(A)** Two methods measuring sequence composition bias are represented for each gene in the *Ddad*3937 genome as follows: GC content at the third codon position index (GC3) in the y-axis, and the Kullback-Leibler distance of dinucleotide frequency distributions (DINT_KL) relative to the *Ddad*3937 genome in the x-axis. The coding sequences from *Ddad*3937 comprise the background set colored in grey. From this set, specific genes are colored according to both (a) the orthologous conservation in SRP genomes and (b) distance to the *Ddad*3937 genome in both GC3 and DINT_KL metrics. Those genes exhibiting above-threshold (mean±SD) values (‘above-threshold’) are represented in light brown if they are genus-specific (orthologs found in Dickeya but not in Pectobacterium genomes). Those genes exhibiting below-threshold values in either GC3 or DINT_KL (‘below-threshold’) are represented in green. Four genes of interest in this study are labelled as follows: *pho*P (*DDA3837_RS11500*), *sly*A (*DDA3937_RS12595*), *exp*R2 (*DDA3937_RS20730*), and *exp*I (*DDA3937_RS20725*). **(B)** The graph follows the same representation described in (A). All the analyses presented above for *Ddad*3937 and the Dickeya genus are mirrored for *Pcb*1692 and the Pectobacterium genus in ‘B’. Different functional classes are labelled as follows: Transcriptional regulators - *pho*P (*PCBA_RS01290*), *sly*A (*PCBA_RS02460*), *exp*R2 (*PCBA_RS20280*), and *exp*I (*PCBA_RS15660*). **(C)** The entire set of genes regulated *in planta* (at early or late infection) by PhoP in *Pcb*1692 comprise the background set colored in grey. Genus-specific genes are highlighted in (a) green if below-threshold, (b) light brown if above-threshold and regulated at early infection, (c) pink if above-threshold and regulated at late infection, or (d) dark brown if above-threshold and regulated at both early and late infection. Genes from different *Pcb*1692 operons are labelled as follows: type III secretion operon - *hrp*A (*PCBA_RS14175*), *hrc*C (*PCBA_RS14210*), *hrp*V (*PCBA_RS14220*), *big*A (*PCBA_RS14230*); carbapenem biosynthesis-associated - *car*R (*PCBA_RS04390*), *car*D (*PCBA_RS04370*), *car*C (*PCBA_RS04375*); and the transcriptional regulator *exp*R1 (*PCBA_RS15665*). **(D)** The graph follows the same representation described in (C) depicting SlyA regulons (early and late infection) in *Pcb*1692. Genes associated with carbohydrate metabolism regulation are labelled as follows: the gene labelled ‘sis’ which encodes a SIS-containing product (*PCBA_RS22170*), *rpi*R (*PCBA_RS22175*), (*PCBA_RS22170*) *deo*R1 (*PCBA_RS02575*).

As for quorum sensing regulators, the *exp*R2/*vir*R homologs showed stronger support for horizontal acquisition in *Ddad*3937 (*DDA3937_RS20730*), supported by both GC3 and DINT_KL, than in *Pcb*1692 (*PCBA_RS20280*), which could indicate that Pectobacterium and Dickeya lineages acquired *exp*R2 independently (Figure 3A and B). This hypothesis will be further discussed in a subsequent section. Moreover, the Pectobacterium-exclusive PhoP-regulated transcriptional regulator *exp*R1 (*PCBA_RS15665*) was strongly supported as a horizontal acquisition by both parametric methods in *Pcb*1692 (Figure 3C). This result is further corroborated by a similar profile observed in the *Pcb*1692 *exp*I gene (*PCBA_RS15660*). Furthermore, the carbapenem biosynthesis transcriptional regulator *car*R (*PCBA_RS04390*) is another GS gene in *Pcb*1692 that has been successfully supported as an HGT candidate with high confidence (Figure 3C). In addition, not only *car*R apparently has been horizontally transferred into the *Pcb*1692 genome, but it is the only gene in the carbapenem biosynthesis gene cluster supported by both parametric methods for HGT prediction (S2 Table). *car*R is also the only GS gene in the *car* gene cluster (S1 Table). These results strongly suggest that *car*R was acquired by *Pcb*1692 independently (probably at a later stage) from *car*A/B/C/D/E/F genes. Also, the presence of two QS regulators (*exp*R1 and *car*R) along with two conspicuous genomic regions encoding bacterial secretion systems (i.e. T3- and T6SS) under PhoP regulation in *Pcb*1692 may be correlated. Hence, we conducted an in-depth investigation of the PhoP-dependent regulation of these secretion systems and the phenotypic outcomes that this regulatory association may give rise to.

### PhoP-dependent regulation of secretion systems and their effects on virulence and fitness

The PhoP-dependent regulation of Pectobacterium-exclusive QS regulators, along with type III and VI secretion systems prompted us to measure both the ability of *Pcb*1692Δ*pho*P to (a) outcompete other SRP *in planta*, and (b) cause tissue maceration in potato tubers. The *Pcb*1692Δ*sly*A was also included for comparative means. Therefore, we first performed *in planta* bacterial competition assays in order to determine the relative contribution of PhoP in bacterial competition. For this experiment, *Ddad*3937 was selected because it is a well-studied niche competitor of *Pcb*1692 in potato tubers. The growth of *Ddad*3937 when co-inoculated *in planta* with *Pcb*1692Δ*pho*P displayed a five-fold reduction (CFU/ml), as opposed to a three-fold reduction when co-inoculated with *Pcb*1692 wild-type (Figure 4A). Conversely, the *Pcb*1692Δ*sly*A strains inhibition against *Ddad*3937 exhibited no significant difference when compared to that of the wild-type *Pcb*1692. The results imply a direct outcome of the overexpression of competition-related mechanisms following *pho*P deletion. Arguably, one of the major players in this increased *in planta* competition aggressiveness might be the extensive array of 24 T6SS-related genes overexpressed in the absence of PhoP.

**Figure 4.**
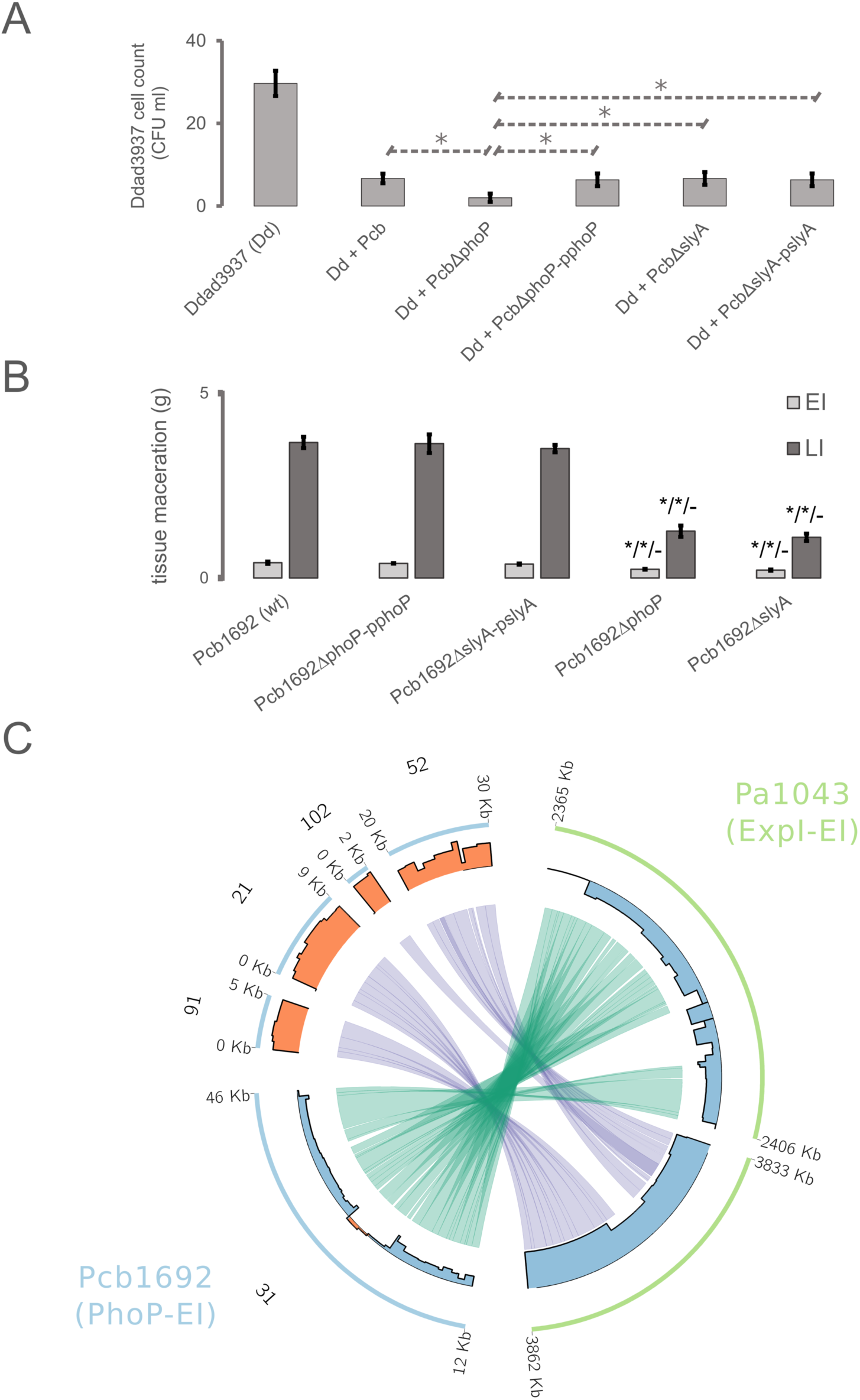
*In planta* bacterial competition and virulence of *Pcb*1692 strains and the transcriptional variation of type III and VI secretion systems in *pho*P and *exp*I mutants respectively in *Pcb*1692 and *Pa*1043. **(A)** The graph depicts the survival of *Dickeya dadantii* strain 3937 (*Ddad*3937) in potato tubers when co-inoculated with different strains of *Pectobacterium carotovorum* subsp. *brasiliense* strain PBR1692 (*Pcb*): *Pcb*Δ*pho*P, *Pcb*Δ*sly*A (mutants) *Pcb* (wild-type), *Pcb*Δ*pho*P*-*p*pho*P, *Pcb*Δ*sly*A*-*p*sly*A (complemented mutant strains) measured in colony-forming units (CFU/ml). Significant differences between different samples were analyzed using Student’s t-test (* for *P* < 0.05). **(B)** The weighed macerated tissue from potato tubers infected with *Pcb*1692 strains was measured and the mass is expressed in grams (g). The strains analyzed are represented as follows: *Pcb*1692 wild-type (wt); *Pcb*1692 *pho*P mutant (*Pcb*1692Δ*pho*P); *Pcb*1692 *sly*A mutant (*Pcb*1692Δ*sly*A); *Pcb*1692 *pho*P mutant complemented with *pho*P (*Pcb*1692Δp*pho*P); *Pcb*1692 *sly*A mutant complemented with *sly*A (*Pcb*1692Δp*sly*A). Above the mutant’s measurement bars, the sequence of three Student’s t-test results is depicted in which the mutants’ macerated masses are compared respectively with that of (i) wild-type, (ii) respective complement and (III) the other mutant (*Pcb*1692Δ*pho*P is compared to *Pcb*1692Δ*sly*A and vice versa). **(C)** Genomic regions from *Pcb*1692 (light-blue) and *P. atrosepticum* strain scri1043 (*Pa*1043) (light-green) are represented in circular ideograms. Coordinates of 5’ and 3’ ends of each genomic region are displayed in kilobases (kb). In the *Pcb*1692 ideograms, respective contig numbers are labelled outside of the ideogram. In the inner radius, the bar plot indicates, for each region, the transcriptional variation level (log2 fold change) found in each transcriptome study (either (21) for *Pa*1043, or this study for *Pcb*1692). The bars are colored to highlight up-regulation (orange) or down-regulation (blue). The inner links binding genomic regions represent the homologous range of type VI (purple) and III (green) secretion systems conserved between *Pcb*1692 and *Pa*1043 genomes.

Next, both *Pcb*1692Δ*sly*A and *Pcb*1692Δ*pho*P strains had their virulence against potato tubers examined by measuring the extent of maceration in early and late infection. The results showed that despite the positive regulation of T6SS- and PCWDE-encoding genes in the *Pcb*1692Δ*pho*P, this strain was attenuated in virulence similar to the *Pcb*1692Δ*sly*A with no statistically significant difference between them (Figure 4B). In comparison to the bacterial competition assay, tissue maceration comprises a complex phenotype that encompasses a wide range of necessary factors. Thus, the transcriptional disruption of a large array of genes in the four regulons (PhoP-, SlyA-EI, PhoP, SlyA-LI), ranging between 66-318 protein-coding genes may be responsible for the attenuated tissue maceration in these mutants (S1 Table).

Aiming to inspect the possible PhoP-regulated mechanisms involved in this increased *in planta* growth inhibition by *Pcb*1692 over *Ddad*3937, we will next lay emphasis on secretion systems regulation. The T6SS is a protein complex assembled through both bacterial membranes featuring a contractile sheath and an injectisome-like structure composed by haemolysin co-regulated proteins (Hcp) that delivers specialized effectors into target cells (31). Both T6 structural- and effector-encoding genes were found under PhoP regulation, totalling 24 T6SS-related genes consistently overexpressed upon *pho*P deletion across the early and late infection (S1.1 and S1.2 Tables). By comparing evidence found from PhoP network in *Pcb*1692 with those from previously reported analyses on a QS-mutant strain of *P. atrosepticum* (*Pa*1043) during infection on potato tubers (21), some important observations were made. For this analysis, the fundamental difference between the two mutant strains, i.e. *Pcb*1692Δ*pho*P and *Pa*1043Δ*exp*I, is that the *exp*R1 regulator is overexpressed in the first and repressed in the second. In the *Pa*1043Δ*exp*I strain, 23 T6SS genes exhibited a strong transcriptional decrease in *Pa*1043 upon the disruption of *exp*I/*exp*R1 (Figure 4C). Contrarily, *Pcb*1692 overexpresses *exp*R1 along with 24 T6SS genes in the absence of *pho*P. This pattern shows a direct interference of QS disruption over the transcriptional modulation of T6SS genes in *Pa*1043. Additionally, it strongly suggests that the PhoP-dependent regulation of the T6SS gene cluster in *Pcb*1692 could primarily depend on the *exp*R1 regulator.

Moreover, we analyzed the T3SS regulation in the two mutants (Figure 4C). Curiously, both *pho*P and *exp*I mutants exhibited a similar pattern of extensive down-regulation of the T3SS genes. In *Pcb*1692, the PhoP transcriptional impact encompasses 23 genes. Of these, 12 have detectable orthologs in *Pa*1043 under the transcriptional influence of QS. In this context, since *exp*R1 homologs are regulated in opposite directions in each mutant, the PhoP network in *Pcb*1692 seemingly regulates several elements from the T3SS in a QS-independent manner.

### Characterizing new gene families within the T3SS operon in Pectobacterium

During the secretion systems regulation analyses, we found two GS PhoP-regulated genes unannotated by BlastP-based methods within a T3SS-encoding cluster in *Pcb*1692 (*PCBA_RS14220*; *PCBA_RS14230*). By examining the sequences from their products, we found no detectable functional domains that could indicate their role. Hence, aiming to classify these unannotated genes, we implemented sensitive sequence comparison techniques (see ‘Methods’ for details) in order to examine their primary structure and search for conserved motifs. This analysis led to the identification of one novel motif in each protein, which are associated respectively to (a) <u>b</u>acterial <u>I</u>g-<u>l</u>ike (Big) and cadherin domains, and (b) hrp-related chaperones.

After extensive iterative searches against the UniprotKB online database via the Hmmer search tool (32), and sequence alignment inspections, we gathered respectively 538 and 429 similar sequence sets to the Big-associated and the hrp-related. The Big-associated motif, found in PCBA_RS14230, exhibits highly conserved Tyr and Leu residues respectively in the second, and prior to the first β-sheets, which may represent core residues in their structure (Figure 5A). This conserved motif was found primarily in sequences exhibiting no companion domains, similar to those from Pectobacterium genomes. Representative genes encoding this exact architecture are most common in Gammaproteobacteria, encompassing diverse lifestyles such as animal and plant pathogens (e.g. Vibrio and Pectobacterium genera), or insect gut symbionts (e.g. Gilliamella genus). Additionally, Alpha- and Betaproteobacteria also carry genes encoding the same architecture, including organisms from Burkholderia and Mesorhizobium genera (Figure 5B). Aside from the instances in which these Big-associated motifs are found solo in a given sequence, they are even more frequently found in long (∼1000 amino acids) proteins constituted by numerous domain repeats. These repeats often include combinations of Cadherin and/or Big domains (Pfam: PF00028; PF02369), along with the novel Big-associated. These families are known for their typical association with phage-like structures and have been linked to carbohydrates recognition in the cell surface enabling phage adsorption (33), which will be further discussed below.

**Figure 5.**
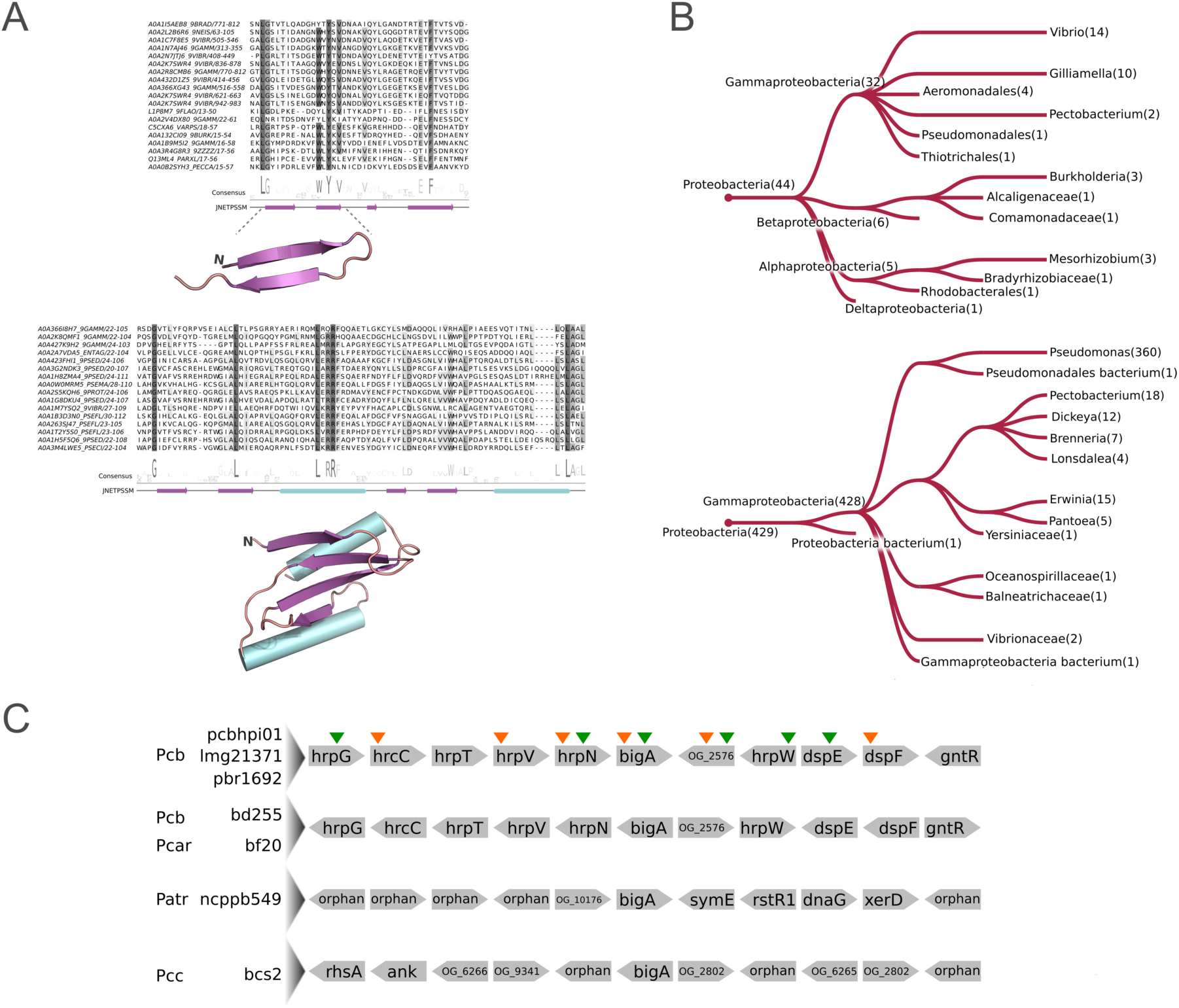
Multiple sequence alignments, structural scaffolds and phyletic distributions of two conserved motifs in type III secretion system-related putative effectors found in *Pcb*1692. **(A)** The degree of conservation of each amino acid column is highlighted in increasingly darker shades of grey. Below each alignment, the amino acids logo depicts the amino acid consensus among the representative sequences. Next, secondary structure predictions are displayed in both linear and folded (obtained through sequence modelling) configurations. Purple arrows represent β-sheets, and cyan cylinders represent α-helices. **(B)** Taxonomic groups in which the Big-associated (top) and HrpV-borne (bottom) motifs are found in sequences exhibiting the same domain architecture as in *Pcb*1692 are depicted in the phylogenetic trees. Taxonomic distributions were assessed through Hmmsearch online tool (32). The branches are labelled with the correspondent number of genes found in the group. **(C)** Genomic architectures of seven Pectobacterium strains are represented depicting five genes up and downstream in *big*A gene neighborhoods. Gene names were annotated according to sequence searches on eggNOG database where applicable. Sequences unannotated by eggNOG are either represented by their respective orthologous group labels (see ‘Methods’ for details) or regarded as ‘orphans’ in case no orthologous sequences were identified across the SRP genomes. Different species are abbreviated on the left as follows: Pcb (*P. carotovorum* subsp. *brasiliense*), Pcar (P. carotovorum), Patr (P. atrosepticum), Pcc (*P. carotovorum* subsp. *carotovorum*). Species names are adjacent to strain names. Arrows above the top architecture indicate up-/down-regulation (up- or down-arrow respectively) of the respective genes by PhoP in *Pcb*1692 (strain pbr1692) either at early infection (orange), or late infection (green) on potato tubers.

In the second unannotated gene product (PCBA_RS14220) we found a conserved α_2_β_4_ structure, which was underpinned by 429 similar sequences gathered during the iterative searches described above (Figure 5B). Approximately 45% of the significant hits found were annotated as HrpV, which form chaperone heterodimers with HrpG to regulate T3SS expression (34). Importantly, besides this newly found α_2_β_4_ motif, no other annotated domains could be detected in any of those 429 HrpV-like sequences. Next, we investigated the structural conservation of those sequences using HHpred (35), which detected similarities with type III chaperones, such as those from SchA, or CesT (Protein data bank: 4G6T, 5Z38). Thus, although the primary structure does not match any currently annotated domain, the α_2_β_4_ fold found in these sequences were recognized as being similar to type III chaperone-related by HHpred. The impossibility of annotating the HrpV sequence via Blastp-based methods may be a consequence of extensive sequence variations occurred in different organisms. This notion is supported by our orthology analysis, which has split the HrpV sequences from Pectobacterium (represented in 50/61 genomes) and Dickeya (represented in 39/39 genomes) into two separate groups (S3 Table). Nonetheless, the hidden Markov model (hmm) profile consolidated in this analysis (see ‘Methods’ for details) was successfully detected in both groups.

Next, we inspected gene neighborhoods of *hrp*V and the Big-associated (*big*A) genes across SRP genomes. As expected, the 89 *hrp*V genes from both orthologs groups, i.e. OG_3591 (Pectobacterium) and OG_3957 (Dickeya), are consistently surrounded by T3SS-related genes (S4 Table). However, this pattern is not observed in *big*A orthologs. Gene neighborhood screening of seven out of ten genomes carrying *big*A orthologs (three genomes were not suitable for the analysis due to incompletion of genome assembly) revealed three different types of neighborhoods. Four *Pcb* strains and one strain of *P. carotovorum* exhibit *big*A homologs within T3SS gene clusters (Figure 5C). In one strain of *P. atrosepticum* the bigA homolog is adjacent to phage elements including homologs of the transcriptional repressor *rst*R1, the DNA primase *dna*G, and the site-specific recombinase *xer*D. The strains of *P. carotovorum subsp. carotovorum* have their bigA homolog surrounded by unannotated genes and ‘orphan’ genes that could not be clustered with any other sequences from SRP during the orthology analysis. Furthermore, the newly identified Big-associated gene b*ig*A described above was predicted as a horizontally transferred gene (Figure 3C). By inspecting the surrounding genes within the *Pcb*1692 T3SS cluster, the *big*A gene is the only that exhibits full support from the HGT prediction (Figure 3C and S2 Table). This result suggests that *big*A acquisition is posterior to the consolidation of the T3SS in the *Pcb*1692 genome, which is consistent with the above genus-specific prediction for *big*A. Together these results elucidated the presence of HrpV and the HGT candidate *big*A under PhoP regulation. Moreover, the *in planta* transcriptional regulation of a known T3SS regulatory element (i.e. *hrp*V) by PhoP points to an additional layer of control over T3SS, which will be addressed in the ‘Discussion’ section.

## DISCUSSION

### The influence of lineage-specific rearrangements in the PhoP and SlyA regulatory networks on host-adaptation

It has been reiterated that the birth of novel functions in prokaryotes is mainly driven by HGT, instead of by duplication and divergence (36, 37). For instance, it has been estimated that in representatives of both *Escherichia* and *Salmonella* genera, more than 98% of protein family expansions arise from HGT (37). Moreover, even gene copy increments in bacterial genomes can often be a consequence of HGT, rather than duplication events (7, 38). As a consequence, the fixation of newly acquired genes requires that existing regulatory circuits reshape accordingly, allowing those new elements to fit properly into specific transcriptional programs. This often results in the recruitment of new genes into existing transcriptional networks (5). As our results showed, PhoP, SlyA, and PecS exhibited a consistent bias towards the regulation of GS-HGT candidate genes at late plant infection. This indicates that transcriptional mobilization of recently acquired genes by these regulators may play a particularly important role in the late stages of infection in these organisms. These findings may even point to a broader trend, in which these distinctive traits are preferentially recruited in late stages of the soft rot disease. At this point in time, with the increasing availability of nutrients in consequence of plant cell lysis (14), the demand for mobilization of distinctive traits in closely related lineages could be specifically focused on interspecies competition.

The participation of PhoP in the regulation of GS genes was also reported in the Salmonella genus, as ∼50% of the genes found under PhoP regulation has no orthologous counterparts outside this group (39). This evidence is in accordance with the overrepresentation of GS genes found under PhoP control in *Pcb*1692. It also reinforces the PhoP tendency towards regulating GS loci, despite the contrasting lifestyles between most Salmonella and Pectobacterium representatives. A similar pattern has been reported in *Yersinia pestis* and *Y. enterocolitica* within the RovA regulatory network (40). That report uncovered horizontally acquired genes within the RovA regulon, including genes associated with disease development (40). The *rovA* gene belongs to the MarR family of regulators, from which the best-characterized members – i.e. RovA and SlyA – have been linked to direct binding site competition with the histone-like nucleoid structuring regulator (H-NS) (41, 42). The H-NS role in the repression of horizontally acquired regions is widely characterized in gram-negative bacteria, as it typically works through the recognition of AT-rich regions (43, 44). These observations converge with the results presented here as 32.8 and 47% of all genes regulated by SlyA respectively at early or late infection were successfully predicted, by at least one parametric method, as HGT candidates.

Of particular importance, our results shed light on the relevance of transcriptional network rearrangement processes for PhoP network expansion, which includes the acquisition of other transcriptional regulators such as *car*R, *exp*R1. The evidence found here for the horizontal acquisition of *car*R and *exp*R1 is in accordance with previous phylogeny-based predictions of *lux*I/*lux*R horizontal transfer within the *Proteobacteria* phylum (45), as well as between gram-negative and -positive bacteria (46). These two regulatory connections, i.e. *pho*P-*car*R and *pho*P-*exp*R, represent an important innovation in the SRP group that impacts the regulation of several host adaptation-related systems in *Pcb*1692, and possibly other species in the Pectobacterium genus.

### *deo*R1 and *rpi*R transcription factors and the prominent role of SlyA in carbohydrate metabolism regulation

As previously highlighted, the transcriptional regulator *deo*R1 homologs spread across 53 out of 61 Pectobacterium genomes, whereas the Dickeya lineage apparently lost this gene. DeoR-type regulators have been recognized for a long time as transcriptional repressors of carbohydrate metabolism-related genes (47, 48). In the gram-positive soil-borne bacterium *Corynebacterium glutamicum*, the DeoR-type regulator SugR was shown to repress the transcription of phosphoenolpyruvate-dependent phosphotransferase system genes, such as *pts*G, *pts*S and *pts*F (49). Also, another DeoR family member termed UlaR was identified as a repressor of the l-ascorbate gene cluster (*ula*) in *E*. *coli*; this repressor was shown to release the *ula* promoter region upon binding to the l-ascorbate-6-phosphate molecule (50). These reports are strikingly supported by our data set, in which upon the overexpression of *deo*R1 in the *sly*A mutant, 20/23 genes annotated in the ‘Carbohydrate metabolism’ KEGG term (09101) are consistently down-regulated at early infection (Figure 6). These observations suggest that SlyA may directly or indirectly repress the *deo*R1 homolog in the wild-type *Pcb*1692, which in turn may release DeoR1 from repressing an array of carbohydrate metabolism-associated genes. Importantly, 69.6% (16/23) of the carbohydrate metabolism-related genes repressed in SlyA-EI, were also repressed in the PhoP-EI regulon (Figure 6 and S1.1 and S1.3 Tables). Thus, since *Pcb*1692Δ*pho*P and *Pcb*1692Δ*sly*A mutant strains exhibit opposite patterns of *sly*A regulation, the evidence indicates that normal expression levels of both *pho*P and *sly*A are required for the expression of these carbohydrate metabolism-associated genes *in planta*.

**Figure 6.**
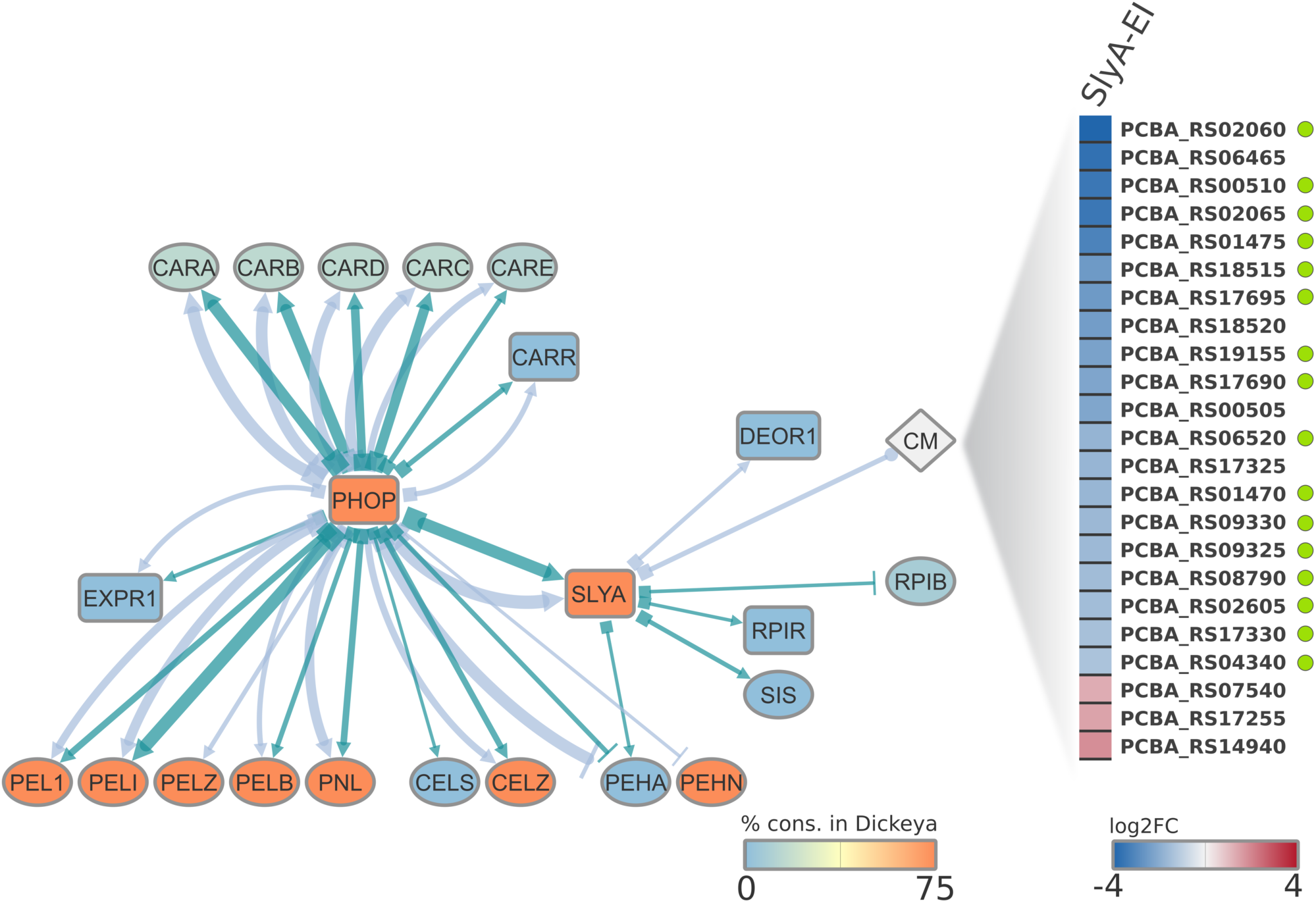
Regulatory interplay between PhoP, SlyA and QS systems in *Pcb*1692. Three subsets of host adaptation and/or virulence themes were extracted from both PhoP and SlyA *in planta* regulons. Transcriptional regulator genes in the network are highlighted in square shapes. The diamond shape represents an array of genes annotated by the KEGG term 09101 ‘Carbohydrate metabolism’ (CM). Each node is colored according to the relative presence in the Dickeya genus according to the color scale at the bottom of the network. The target link shape depicts the link relationship between source and target nodes: up-regulation (arrow), down-regulation (perpendicular line), or mixed (circular shape). The source link shape is represented as square to indicate “in the absence of”, which is the direct inference based on the RNA-Seq experiment performed using the two mutants (*Pcb*1692Δ*pho*P and *Pcb*1692Δ*sly*A). Increasingly thicker links in the network represent higher |log2 fold change| values. Light-blue and dark-green links respectively indicate regulation at early and late infection by one of the two analyzed regulators (PhoP or SlyA). On the right side of the network, a one-column heatmap depicts the differential expression (log2 fold change) in genes associated with carbohydrate metabolism regulated by SlyA at early infection. Genes are represented by the respective locus tags according to the NCBI database. Green circles adjacent to individual genes highlight those for which PhoP-dependent regulation at early infection was also found.

Converging with this observation, the GS-HGT candidate *PCBA_RS22175* seem to encode a RpiR transcriptional regulator homolog. This gene is regulated by SlyA exclusively at late infection, indicating a particular demand for the transcription of this recently acquired *rpi*R homolog at this stage. The *rpi*R gene family is frequently associated with carbohydrate utilization regulation (51, 52). Specifically, the original characterization of the *rpi*R gene in *E. coli* by Sorensen and Hove-Jensen (53) revealed its repressive effect over the ribose phosphate isomerase B (*rpi*B) transcription. The *rpi*B interconverts ribulose 5-phosphate and ribose 5-phosphate and is necessary for the utilization of ribose as a carbon source (54). In our data set, this notion is supported by the down-regulation of *rpi*B and concomitant up-regulation of *rpi*R in the SlyA late infection regulon *in planta* (Figure 6 and S1.4 Table). Besides *rpi*R, another HGT candidate gene encoding the SIS domain (Pfam: PF01380) was found in the SlyA-LI network (*PCBA_RS22170*). The SIS domains can be found solo in protein sequences, such as in GmhA (55), or accompanied by other domains (56). Interestingly, the *Pcb*1692 gene *PCBA_RS22170* encodes a solo SIS domain, whereas its neighbour *rpi*R homolog *PCBA_RS22175* encodes a solo HTH_6 (Pfam: PF01418). The fact that in 95% (2542/2677) of the *rpi*R homologs (eggNOG: COG1737) (57) contain both the DNA-binding domain HTH_6 and the SIS domain, suggests that these two entries (i.e. *PCBA_RS22170* and *PCBA_RS22175*) may be in fact a single gene in *Pcb*1692. Together, the evidence sustains that SlyA acquired a novel regulatory system that controls an important step of the pentose phosphate pathway through network rearrangement.

### PhoP/QS interplay *in planta* in *Pcb*1692: Transcriptional regulation of carbapenem biosynthesis and PCWDE encoding genes

Two out of three quorum sensing regulator homologs from the *Pcb*1692 genome were found under PhoP transcriptional regulation: *expR*1 and *carR*. The absent *expR* homolog in the PhoP regulons is the *expR2*/*virR* gene, which has been shown to regulate several virulence themes, such as iron uptake, motility, and expression of PCWDEs in *P. atrosepticum* (58, 59). The mechanistic synergy between ExpR1 and 2 has been previously described by Sjoblom, Brader (29), in which both regulators are implicated in the control of virulence-related genes in Pectobacterium. In that study, they also observed ExpR2 capability of sensing a broader range of autoinducer (AI) molecules when compared to ExpR1 (29). Notably, QS has been previously reported to be controlled by other global regulators. For example, both in *Pseudomonas syringae* (60) and in *Pcb*1692 (61), the relationship between the ferric-uptake regulator (Fur) and the QS system was determined through contrasting concentrations of N-acylhomoserine-lactone produced by the *fur* mutant and wild-type strains. Also, the regulation of QS by PhoQ/PhoP observed in *P. fluorescens* seems to be affected by the Mg^2+^ concentration level in the medium (62).

The biosynthesis of the carbapenem antibiotic is one of the important QS-subordinate systems. CarR-dependent QS regulation in *Pectobacterium carotovorum* subsp. *carotovorum* relies on the presence of N-(3-oxohexanoyl)-L-homoserine lactone (OHHL) ligand (28). Interestingly, the stability of both carbapenem and the OHHL molecules are affected by pH variations (63, 64). In this context, the PhoPQ two-component system ability to respond to pH variations (65, 66) may (at least partially) explain the success of recruiting *carR* and other *car* genes into the PhoP regulatory network. Thus, in the absence of *phoP*, *carR* and five other *car* genes are overexpressed *in planta*, which by inference, means that PhoP network suppresses their transcription *in planta* in the wild-type. Hence, PhoP activation, arguably as a consequence of milieu acidification, is able to prevent carbapenem biosynthesis under pH conditions that are unfavourable for the antibiotic stability (Figure 6).

A similar logic might as well apply to the PhoP regulation of *expR*1 and several PCWDEs. Indeed, we observed a marked contrast in PhoP regulation pattern over distinct PCWDE classes exhibiting (a) neutral/alkaline optimum pH (∼6.8-8) such as pectate/pectin lyases and cellulases (19, 20), and (b) acidic optimum pH (∼6) such as polygalacturonases (Figure 6) (67). This regulatory pattern does not correspond to the unidirectional regulation of PCWDE-encoding genes through the QS-RsmA system, which has been repeatedly reported in the Pectobacterium genus (16, 27). Thus, the fact that PhoP network coerces the expression of different sets of PCWDEs in opposite directions at the same stage implies that the QS-RsmA cascade is not the only regulatory mechanism controlling those genes in *Pcb*1692. Therefore, although at first glance our results suggest a direct PhoP-ExpR1-RsmA-PCWDE regulatory hierarchy, an additional fine-tuning step enforced either directly or indirectly by PhoP seems to provide an alternative control system over PCWDE transcription. Moreover, this alternative pathway seems to be RsmA-independent, which follows a previously proposed model of QS regulation (29), since neither *rsm*B or *rsm*A are influenced by PhoP absence in *Pcb*1692. In this context, the regulation of PCWDE by PhoP in *Dickeya dadantii* has been previously reported. However, PhoP seemed to cause a unidirectional impact over pectate lyases and polygalacturonases (4) conversely to what was observed in *Pcb*1692. Such contrast should not be surprising, since the Dickeya spp. lacks the *exp*R1 QS element, which may be an important part of the PhoP-dependent regulation of PCWDE in *Pcb*1692. Thus, the different patterns found in these two genera may occur as a result of the PhoP-ExpR1 interplay in the regulation of PCWDE that seems to take place in Pectobacterium and not in Dickeya spp.

Intriguingly, gene-neighborhood analyses revealed that the *exp*R1 position in Pectobacterium genomes is locked upstream of *exp*I, which is the exact same pattern observed in the *exp*R2/*vir*R homologs across the Dickeya genus (Figure 7A). In terms of conservation, the *exp*R2/*vir*R can be found in 100% of the SRP strains analyzed, and in Pectobacterium genomes, it is mostly surrounded by membrane transport systems and other transcriptional regulators. Whereas the third homolog *car*R, exists only in 21 out of 61 Pectobacterium strains analyzed, and it is consistently adjacent to the carbapenem biosynthetic cluster in the genus, as previously reported by Shyntum, Nkomo (68) (Figure 7A). This agrees with a recent estimate that ∼76% of ExpR/LuxR genes exist in genomes without an adjacent ExpI/LuxI homolog (25). Also, according to the HGT predictions, the acquisition of *exp*R1 and *car*R by *Pcb*1692 appears to have occurred through horizontal transfer in the Pectobacterium lineage. Although some species such as *P. parmentieri* and *P. wasabiae* lost the entire carbapenem biosynthesis cluster as previously communicated by Shyntum, Nkomo (68), the *exp*R1 remains prevalent in the Pectobacterium genus. Thus, the evidence suggests that each of the three *exp*R/*lux*R homologs found in Dickeya or Pectobacterium genera have been acquired horizontally in an independent fashion. Furthermore, horizontal acquisition of *exp*R/*lux*R homologs has been predicted to occur either as a regulatory cassette that includes *exp*I/*lux*I, or individually (45, 69). This notion converges with the results observed in (a) *exp*R2/*vir*R or *car*R found in Pectobacterium genomes, in which individual acquisition is the most parsimonious assumption, and (b) *exp*R2/*vir*R in Dickeya or *exp*R1 in Pectobacterium genomes, which most likely were acquired as a regulatory cassette (Figure 7A).

**Figure 7.**
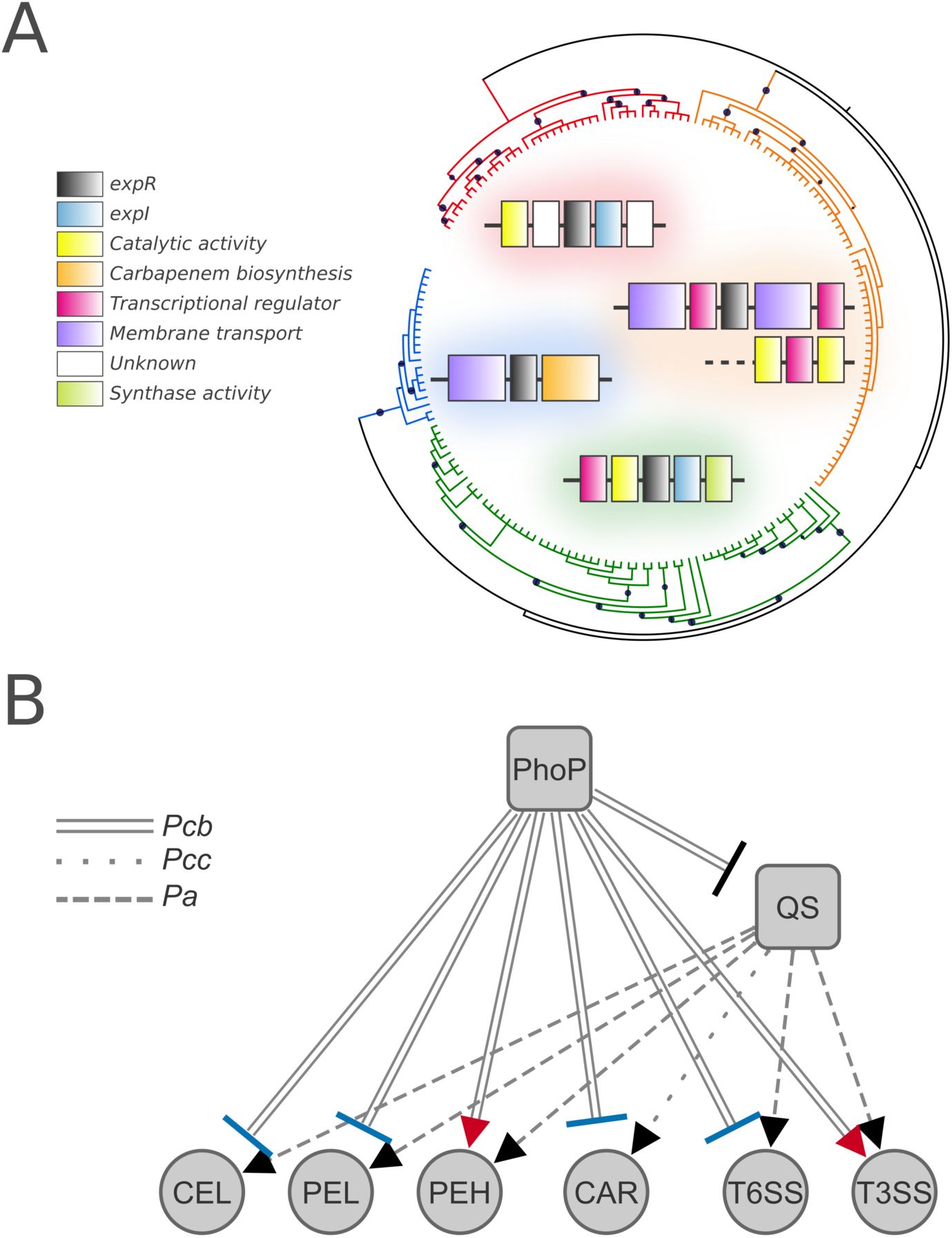
Evolution of ExpR sub-families in distinct genomic contexts in SRP and the summary of PhoP-QS interplay in *Pcb*1692. **(A)** The phylogenetic reconstruction of the autoinducer recognition module found in the ExpR/LuxR homologs from Pectobacterium and Dickeya was inferred by approximately-maximum-likelihood. The tree is represented in unscaled branches with bootstrap values displayed as black circles. The different ExpR/LuxR orthologous groups previously predicted through OrthoMCL, all of which were clustered exclusively with sequences from the same genus, are highlighted in colors as follows: ExpR2/VirR from Dickeya (red) and Pectobacterium (orange) genera; ExpR1 (green) and CarR (blue) from Pectobacterium. For each group, the dominant gene-neighborhoods found across the genus is depicted with the corresponding clade color in the background. **(B)** The summary network is displayed based on indirect inference from the RNA-Seq: if a given gene/system is repressed in the absence of PhoP, then this gene/system is activated in the presence of PhoP. Blue and red target shapes in the network links respectively represent correspondence and conflict between (i) the observed regulation by PhoP in *Pcb*1692 and (ii) the quorum sensing (QS) regulation *in planta* reported in other Pectobacterium species. Regulated pathogenicity themes are abbreviated as follows: cellulases (CEL), pectate/pectin lyases (PEL), polygalacturonases (PEH), carbapenem biosynthesis (CAR), type VI and III secretion systems (T6SS and T3SS respectively), quorum sensing regulators (QS). Link patterns represent the species from which the inference is based on *Pectobacterium carotovorum* subsp. *carotovorum* (Pcc), *Pectobacterium atrosepticum* (Pba) and *Pcb*1692 (Pcb).

Based on previous reports on how QS regulators modulate either PCWDE (21, 70) or carbapenem biosynthesis (28, 71) in SRPs, the possible PhoP-QS interplay can be inferred in our results. Hence, for pectate lyases (Pel), cellulases (Cel) and the *car* genes, the observed regulation of QS by PhoP agrees with those from previous reports, which reinforces the idea of QS regulatory mediation for those genes (Figure 7B). Conversely, since two polygalacturonase-encoding genes exhibit the opposite transcriptional behavior, they may be controlled by a QS-independent mechanism under the PhoP network. This raises the possibility of a cascade within the PhoP network providing additional pH-dependent control over the transcription of some PCWDE, including Pectobacterium-exclusive PCWDEs, which seems to override QS regulation.

### The PhoP interplay with QS in the transcriptional regulation of T3SS and T6SS

The role of T3SS in pathogenicity has been widely explored in a range of taxa, including some from the SRP group. In *Pectobacterium atrosepticum*, a significant reduction in virulence towards potatoes was observed in mutant strains lacking both T3SS structural genes *hrc*C and *hrc*V (72). In that same study, Holeva, Bell (72) also observed that upon the deletion of either the T3SS-related effector *dsp*E or the helper gene *hrp*N, virulence was also reduced. Among the 14 T3SS-related genes found under PhoP network control in *Pcb*1692 at early infection, *hrc*C, *hrc*V, and *hrp*N are present (S1.1 Table). This correspondence implies that the results observed in our virulence assays, in which *Pcb*1692Δ*pho*P exhibited attenuated virulence compared to the wild-type, was (at least partially) impacted by the repression of those T3SS-related genes.

The transcriptional regulation of T3SS involves complex coordination by several regulatory systems, and some have been reported in different bacterial lineages. This includes the QS system (21), as well as other regulators such as SlyA (73), PhoP (74), PecS (9), and Fur (75). Specifically in *Salmonella enterica*, the T3SS termed Spi/Ssa was observably under indirect PhoP control through the SsrB/SpiR system (74, 76). Otherwise, within the SRP group, previous efforts have concluded that PhoP deletion in *Ddad*3937 has no impact over the regulation of T3SS under *in vitro* conditions (77). In contrast with this observation, our results show a wide impact exerted *in planta* by PhoP on T3SS transcription. These contrasting results should be expected since *in vitro* and *in planta* observations present major differences. This contrast indicates that PhoP regulation over T3SS transcription might depend on specific cues that cannot be met *in vitro*. However, since PhoP-T3SS regulatory link has not been assessed *in planta* in *Ddad*3937 so far, which would allow a direct comparison with the results observed in *Pcb*1692, there is not enough data currently available to predict if: (a) The PhoP-dependent regulation of T3SS *in planta* is orthologous among *Pcb*1692 and *Ddad*3937, or (b) if this mechanism evolved at some point within the Pectobacterium lineage. Nonetheless, the PhoP control over T3SS transcription *in planta* observed in *Pcb*1692 had not been reported in phytopathogens so far. Moreover, our SlyA regulon analysis also shows that contrarily to what was reported in *Ddad*3937 by Zou, Zeng (73), *Pcb*1692 does not rely on SlyA to control T3SS transcription.

Unlike T3SS, the T6SS role in plant colonization processes is not deeply understood in phytopathogens and specifically within the SRP group. Yet, it has been reported that T6SS, similarly to T3SS, is under the regulatory control of QS, as observed in *Pseudomonas aeruginosa* (78), and in the *P. atrosepticum* mutant strain lacking the *exp*I gene (21). The T6SS has been functionally implicated in both host defense manipulation and interbacterial competition. However, these two functionalities are not characteristic of any of the five T6SS phylogenetic clades recently reported, and thus cannot be used to separate T6SS phylogenetic groups (79). As an example, the ability of a T6SS-related Hcp protein to facilitate tumorigenesis in potatoes has been reported in the phytopathogenic *Agrobacterium tumefaciens* DC58 (80). In the same species, the homologous T6SS was later implicated in intra- and interspecies competition *in planta* (81). Furthermore, T6SS has been specifically reported as an decisive asset in interspecies competition *in planta* for *Pcb*1692 (68). In agreement with previous reports, our results from *in planta* bacterial competition indicate that the increased transcription of T6SS in consequence of *pho*P deletion in *Pcb*1692 may be an important factor to boost competitive advantage against closely related competitors (Figure 4A and C).

The observed influence of QS over T6SS transcription in *P. atrosepticum* spans through 23 genes, which exhibited decreased transcription intensity upon QS disruption (21). Similarly, T3SS transcription was also impacted by the disruption of QS, as 21 genes were consistently repressed (21). We verified that in the absence of PhoP, the QS receptor *exp*R1 is overexpressed, and so are several T6SS-related genes, which strongly suggests that QS mediates the PhoP regulation over T6SS during infection. Indeed, this hypothesis is supported by Liu, Coulthurst (21) results in *P. atrosepticum*, in which T6SS transcription decreases as the QS system is disrupted (Figure 7B). On the other hand, since T3SS exhibits the opposite regulatory pattern, similar to the one observed in *P. atrosepticum* following QS disruption, it is likely that PhoP does not depend on QS to regulate the T3SS. Hence, PhoP must be able to override QS regulation of T3SS-encoding genes and control their expression. This could result either from PhoP directly binding to their promoter regions or through an alternative QS-independent regulatory cascade.

### New T3SS-related families and the transcriptional regulation of the *hrp* gene cluster by PhoP

The bacterial Ig-like domains are highly promiscuous entities that have been described in structures of (a) adhesins, such as invasins and intimins (82, 83), (b) phage-tail proteins (84), and (c) bacterial surface glycohydrolases (85). The presence of Big-encoding genes in bacteria has been associated with horizontal transfers as a result of its frequent presence in phage genomes (85), which was strongly corroborated by our HGT prediction. Although the precise function of Big domains remains elusive, several lines of evidence point to a role in surface carbohydrates recognition (84, 85). Indeed, from the “guilt by association” standpoint, this could be the case for these newly found BigA proteins, since they are located in a region where T3SS-related products are typically encoded. Also, the analyses conducted in this study support the posterior horizontal acquisition of the bigA gene in the *Pcb*1692 independently from the rest of the T3SS gene cluster.

The other newly found motif in T3SS-related proteins from *Pcb*1692 is comprised of an α_2_β_4_ structure which was found to be similar to that of bacterial HrpV. Curiously, this motif has been identified before in *Erwinia amylovora*, although it has not been deposited in any public domain databases (34). HrpV is known as a negative regulator of the *hrp*/*hrc* gene cluster in *P. syringae* (86). One of the important protein interactions involving HrpV is the ability to bind HrpS. This interaction forms the HrpV-HrpS complex, which blocks the formation of HrpR-HrpS heterohexamers, subsequently hindering HrpS ability to bind and activate the *hrp*/*hrc* promoter (87, 88). This negative regulation imposed by HrpV was shown to be attenuated by its interaction with HrpG, which then generates a double-negative regulatory circuit controlling the expression of T3SS elements (89). We also found that none of the other transcriptional regulators of the T3SS region (i.e. *hrp*G, *hrp*S, *hrp*R, and *hrp*L) are affected by *pho*P deletion, and yet 23 *hrp*/*hrc* genes exhibit differential expression in the PhoP mutant strain. Thus, the expected result of having the main repressor of *hrp*/*hrc* (i.e. *hrp*V) transcription repressed, should be the increased expression of several genes in this region. However, the opposite was observed, as 23 T3SS genes are consistently repressed across early and late infection stages. These observations imply that an alternative regulatory mechanism employed by the PhoP network is able to coordinate the expression of those 23 *hrp*/*hrc* genes, one that is independent of HrpG/S/R/L. As previously discussed, QS does not seem to be the mechanism responsible for this regulation, based on the results from PhoP regulon analyses combined with past observations (21). This means that PhoP is either directly or indirectly involved in the regulation of T3SS, apparently independently from Hrp-borne regulators or the QS system. Indeed, this hypothesis adds complexity to the topic if taken together with the conclusions reported by Bijlsma and Groisman (74) in *Salmonella enterica*, in which PhoP is responsible for the post-transcriptional regulation of a T3SS gene cluster.

By elucidating the mechanisms of network expansion through transcriptional rearrangement in an important phytopathosystem, we found a wide range of host adaptation- and environmental fitness-related traits being co-opted by SlyA and especially by PhoP. Here we uncover the success of the centralizing strategy that recruited carbohydrate metabolism regulation along with virulence-related systems controlled by QS in *Pcb*1692 into the stress-response regulator PhoP. This seems to provide optimized control over specific systems that may be sensitive to certain environmental conditions that can be recognized by PhoQ/PhoP two-component system.

## MATERIAL AND METHODS

### Growth conditions and construction of *Pectobacterium carotovorum* subsp. *brasiliense* PBR1692 mutant strains

The strains and plasmids used are listed in Table 2. Luria-Bertani (LB) broth and agar plates were used for growing all bacterial strains at 37°C. Different antibiotics were used to supplement the media: kanamycin (50µg/ml), ampicillin (100µg/ml) (Sigma Aldrich) when needed. All reagents were used according to the manufacturer’s instructions. Then, in order to generate the mutant strains, the ASAP database BLASTN tool was used to identify the *Pcb*1692 *pho*P and *sly*A genes (*PCBA_RS01290* and *PCBA_RS02460*). Both *Pcb*1692Δ*pho*P and *Pcb*1692Δ*sly*A mutants were generated using a strategy developed by Datsenko and Wanner (90). Briefly, the up and downstream regions flanking the target genes (approx.1000bp) were amplified using specific primers (Table 3) using polymerase chain reaction (PCR). Kanamycin resistance gene cassette was amplified from pKD4 plasmid. The resulting upstream, kanamycin cassette and the downstream fragments were fused using primers denoted in Table 3 as previously described by Shyntum, Nkomo (68). The fused PCR product was purified and electroporated into electrocompetent *Pcb*1692 cells with pkD20. The resulting transformants were selected on nutrient agar supplemented with 50µg/ml kanamycin. HiFi HotStart PCR Kit (KAPA system) was used in all PCR reactions. The PCR conditions were set as follows: initial denaturation at 95°C for 3 min, followed by 25 cycles of denaturing at 98°C for 30s, annealing at 60-65°C (depending on the primer set), extension at 72°C for 2 min and a final extension at 72° for 2 min. The *Pcb*1692Δ*pho*P and *Pcb*1692Δ*sly*A mutant strains was verified by PCR analyses and nucleotide sequencing.

**Table 2.** List of bacterial strains and plasmids used in this study.

| Bacterial strains | Description | Sources |
| --- | --- | --- |
| *Pectobacterium carotovorum* subsp. *brasiliense* PBR1692 (*Pcb1692*) | Initially isolated from potato in Brazil, sequenced strain | Professor A. Charkowski, Wisconsin University (91) |
| *Pcb1692ΔphoP*-*pphoP* | *Pcb1692ΔphoP* expressing the *phoP* gene from the p-JET-T plasmid; Amp<sup>r</sup> | This study |
| *Pcb1692ΔphoP* | *P. carotovorum* ssp. *brasiliense* 1692*phoP*, Kan<sup>r</sup> | This study |
| *Pcb1692ΔslyA* | *P. carotovorum* ssp. *brasiliense* 1692*slyA*, Kan<sup>r</sup> | This study |
| *Pcb1692ΔslyA*-*pslyA* | *Pcb1692ΔslyA* expressing the *slyA* gene from the p-JET-T plasmid; Amp<sup>r</sup> | This study |
| *Dickeya dadantii* 3937 | *Pelargonium capitatum*, Comoros, sequenced | (92) |
| **Plasmids** |  |  |
| pKD4 | Plasmid containing a Kan<sup>r</sup> cassette | (90) |
| pJET1.2/blunt | Bacterial cloning vector containing the *phoP* gene insert, Amp<sup>r</sup> | Thermofischer |
|  | Bacterial cloning vector containing the *slyA* gene insert, Amp<sup>r</sup> |  |
| pJET-*phoP* | Bacterial expression vector expressing *phoP* gene, Amp<sup>r</sup> | This study |
| pJET-*slyA* | Bacterial expression vector expressing *slyA* gene, Amp<sup>r</sup> | This study |
| pKD20 | Temperature sensitive replication ori (repA101ts); encodes lambda Red genes (exo, bet, gam) | (90) |

**Table 3:**
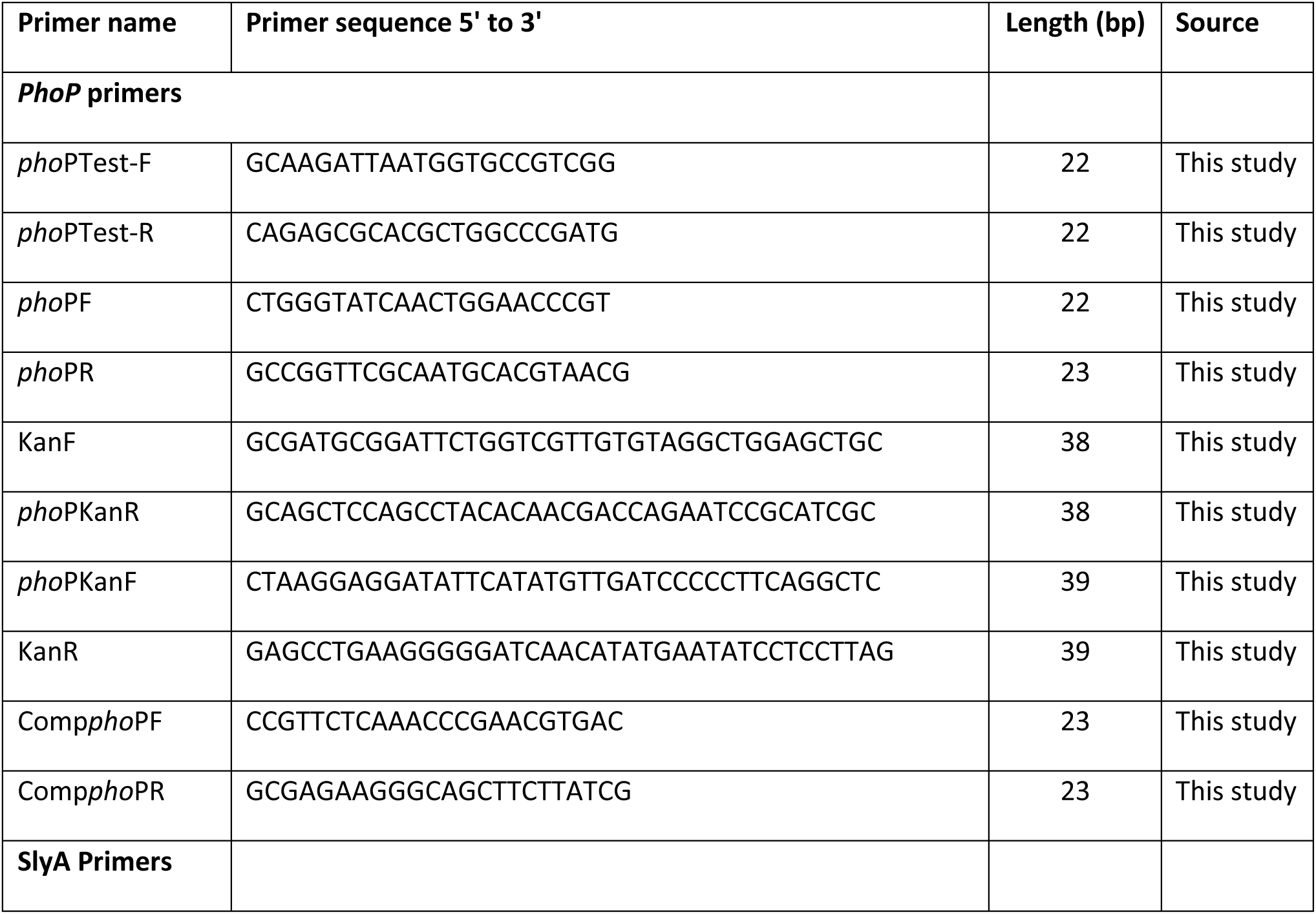
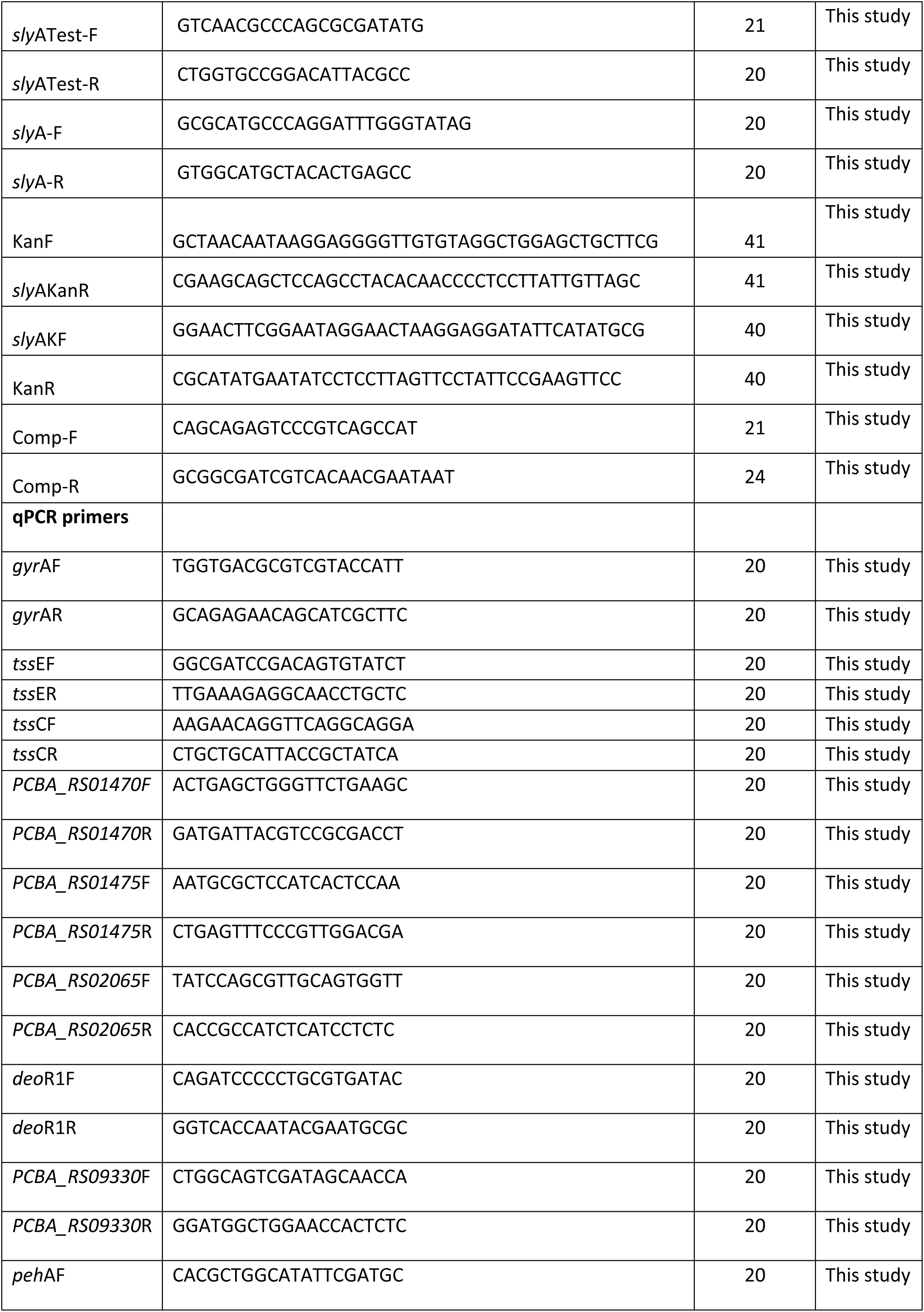
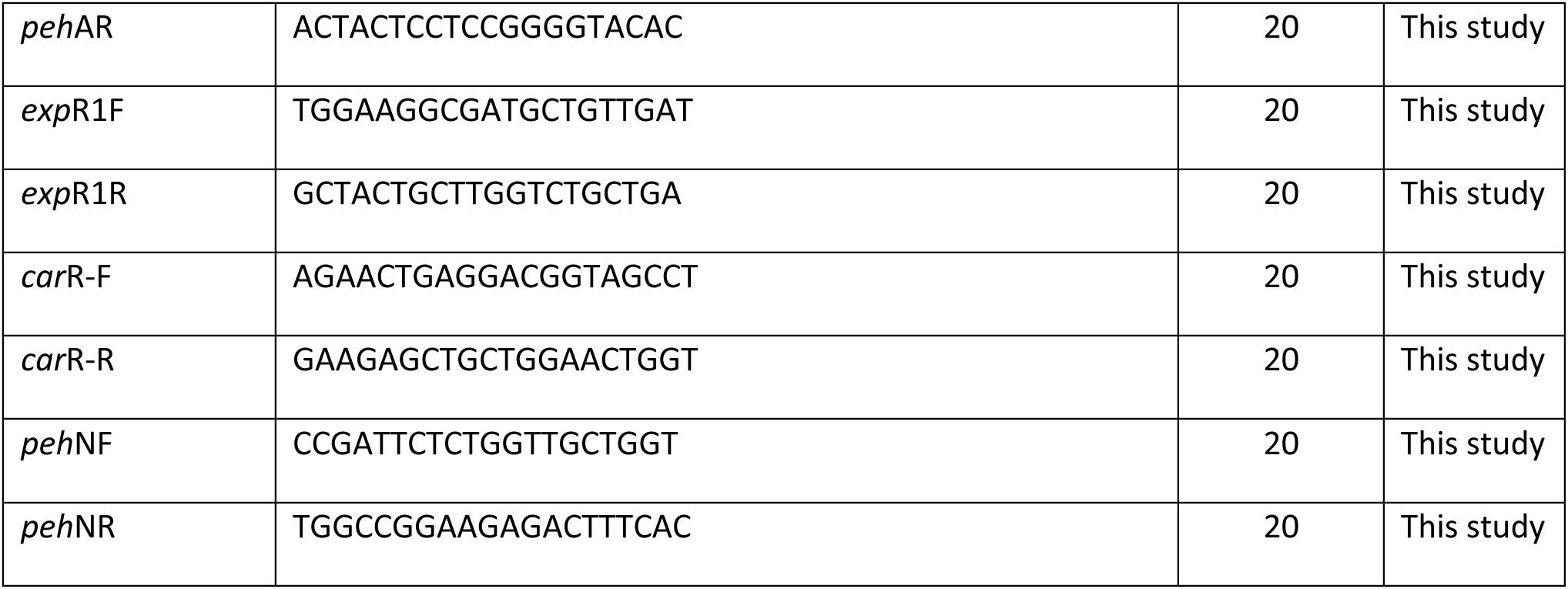
PCR Primers used in this study.

### Complementation of *pho*P and *sly*A mutants

The p-JET1.2/blunt cloning vector was used for the complementation of mutant strains. The *pho*P and *sly*A genes from *Pcb*1692 including ∼500 nucleotides downstream sequence containing the putative promoter sequence were amplified by PCR using complementation primers listed in Table 3. The corresponding fragments were excised from an agarose gel and purified using the Thermo Scientific Gel Extraction Kit according to the manufacturer’s instructions. The fragments were each cloned into p-JET-T to generate pJET-*pho*P and pJET-*sly*A (Table 2). These plasmids were electroporated into electrocompetent *Pcb*1692Δ*pho*P and *Pcb*1692Δ*sly*A mutant strains, transformants (*Pcb*1692Δ*pho*P-p*pho*P; *Pcb*1692Δ*sly*A-p*sly*A) selected on agar plates supplemented with 100 µg/ml Ampicillin and the cloned *pho*P and *sly*A confirmed using PCR and sequencing.

### Tissue maceration assay and total RNA extraction from potato tubers

*Solanum tuberosum* (cv. Mondial, a susceptible cultivar) potato tubers were sterilized with 10% sodium hypochlorite, rinsed twice with double distilled water, air-dried and then stabbed with a sterile pipette to 1cm depth. A 10-μl aliquot of the bacterial cells with OD_600_ equivalent to 1 (*Pcb*1692 wild-type, *Pcb*1692Δ*pho*P, and *Pcb*1692Δ*pho*P*-*p*pho*P) were pipetted into generated holes. As a control, 10 mM MgSO_4_ was inoculated into potato tubers. Holes were sealed with petroleum jelly. Potato tubers were then placed in moist plastic containers and incubated for 12 and 24 hours at 25°C. The macerated tissue was scooped and weighed 12 and 24h post-inoculation to quantify the extent of tuber maceration caused by (*Pcb*1692 wild-type, *Pcb*1692Δ*pho*P and *Pcb*1692Δ*sly*A mutant, and *Pcb*1692Δ*pho*P*-*p*pho*P and *Pcb*1692Δ*sly*A*-*p*sly*A complemented mutant). This experiment was performed in triplicates, three independent times.

For RNA extraction, potato tubers inoculated with *Pcb*1692 wild-type, *Pcb*1692Δ*pho*P, and *Pcb*1692Δ*sly*A were incubated at 25°C with high humidity in plastic containers for 12 and 24 hours maximum. Sampling was done at 12 and 24 hpi by scooping out macerated tissue. Macerated potato tissue from each inoculated site was scooped out and homogenized in double-distilled water. Bacterial cells were recovered by grinding the scooped macerated potato tissues in 20 ml of double-distilled water using autoclaved pestle and mortar. Starch material was removed by centrifuging the ground tissues at 10000 rpm for 1 minute. The supernatant was removed and transferred into new sterilized 50ml Falcon tubes containing RNA stabilization buffer (Qiagen, Hilden, Germany). The experiments were performed using three biological replicates, with three tubers per replicate.

### Determination of total RNA quality

The concentration and purity of each extracted total RNA sample were evaluated using spectrophotometric analysis (NanoDrop® ND-1000; NanoDrop® Technologies, Wilmington, DE) at a ratio of 230/260 nm. The integrity of the RNA was further confirmed using 1% (w/v) agarose gel electrophoresis using 1% TAE buffer at 100 volts for 30 minutes, visualized and image developed using Gel Doc EZ system (Bio-Rad Laboratories, Berkeley, California, USA). Using Agilent 2100 Bioanalyzer (Agilent Technologies, Inc.), total RNA samples’ concentration, RIN and 28S/18S ratio were determined.

### RNA sequencing, Reads mapping, differential expression analysis and genome-wide functional annotation

RNA samples were sequenced in the Novogene facility (823 Anchorage Place, Chula Vista, CA 91914, USA) using an Illumina NovaSeq 6000 machine. The raw data sets are publicly available on the Sequence Read Archive (https://www.ncbi.nlm.nih.gov/sra) hosted by NCBI under the accession number PRJNA565562. Reads sequencing quality analysis was carried out by utilizing *fastqc* software (https://www.bioinformatics.babraham.ac.uk/projects/fastqc). Low-quality segments were then trimmed by Trimmomatic v 0.36 (93). Trimmed reads were next supplied to *hisat2* v 2.1.0 (94) that performed reads alignment to the reference genome of *Pcb*1692 (GCF_000173135.1) obtained from the NCBI database (https://www.ncbi.nlm.nih.gov). The number of aligned reads was subsequently computed by *featureCounts* package (95) and differential expression was analyzed by utilizing the EdgeR package (96), both within the R environment (https://www.r-project.org/). Genes exhibiting transcriptional variation of log2 fold change > 1 (up-regulation), or < −1 (down-regulation), with FDR < 0.05 were assigned as differentially expressed. Mutant samples relative to the wild-type comparisons were done (wild-type *Pcb*1692 X *Pcb*1692Δ*pho*P at 12 and 24 hpi*. Pcb*1692) sequences were functionally annotated by using the eggNOG-mapper tool (97). The annotation provided by eggNOG was next used to retrieve higher annotation levels in the KEGG-library hierarchy (KEGG B and A) (98) through custom scripts written in Perl language (https://www.perl.org/). Enrichment of KEGG terms was determined by one-tailed Fisher exact test and subsequent p-value adjustment by FDR (q-value < 0.05) was performed by custom R scripts using Pcb1692 genome as background dataset. An additional level of annotation was provided by conserved domains inspection on the protein sequences carried out using HMMER3 (99) and the Pfam database (100). Venn diagrams were created using InteractiVenn online tool (101).

### Genus-specific contents, enrichment in specific regulons and genome simulations

The framework used to identify orthologs among strains from Pectobacterium and Dickeya spp. genomes was previously described (6). Briefly, genomic and proteomic information were acquired from the NCBI online database relative respectively to 61 and 39 Pectobacterium and Dickeya strains. All protein sequences were clustered by implementing the OrthoMCL pipeline (102). Next, the presence of representatives from each genus was analyzed throughout the 10 635 orthologous groups using custom Perl scripts. Those gene products that (a) belong to clusters populated exclusively with sequences from one of the genera, or (b) orphans (not clustered), are then considered genus-specific. In order to evaluate the possibility of overrepresentation of GS genes within individual regulons, we used both statistical analysis and computational simulations. Statistical verification was performed in R environment (https://www.r-project.org/) was based on a two-tailed Fisher exact test to assess the correlation between GS occurrence in a given regulon, using the respective strain genome as background. Further, the computational simulations were performed in order to generate 10 000 shuffled copies of the respective genomes (*Pcb*1692 or *Ddad*3937 pseudo-genomes) for each regulon comparison. These comparisons were conducted as follows: for a single regulon, gene positions relative to each regulated gene were retrieved. Next, these exact positions were checked in each of the 10 000 pseudo-genomes for the occurrence of GS genes. The total number of GS genes found in this assessment for each pseudo-genome is computed and compared to the real data.

### Gene expression analysis through qRT-PCR

To validate the differential expression analysis of genes from the RNA-seq data, a qRT-PCR analysis was performed using 6 randomly selected genes. RNA samples used in the qRT-PCR analysis were the same as those used in RNA sequencing and were prepared using RN*easy* mini kit (Qiagen, Hilden, Germany) according to the manufacturer’s instructions. The first-strand cDNA was reverse transcribed from 5 μg of total RNA with the SuperScript IV First-Strand Synthesis System (Invitrogen). Quantitative real-time PCR (qRT-PCR) was performed using QuantStudio™ 12K Flex Real-Time PCR System (Applied Biosystems™). Primers used in this study are listed in (Table 2), the following cycling parameters were used: 95°C for 3 min followed by 40 cycles of 95°C for 60s, 55°C for 45s and 72°C for 60s, followed by 72°C for 7 min; *ffh* and *gyr*A (house-keeping gene control) were used to normalize gene expression. The gene expression levels of the following genes were analysed: *peh*A (*PCBA_RS10070*), *peh*N (*PCBA_RS15410*), *exp*R1 (*PCBA_RS15665*), *car*R (*PCBA_RS04390*), deoR1 (*PCBA_RS02575*), *tss*E (*PCBA_RS11165*), *tss*C (*PCBA_RS11175*), and four PTS permeases (*PCBA_RS01475; PCBA_RS01470; PCBA_RS02065; 09330*). The same method was applied to measure the expression levels of *pho*P gene in *Pcb*1692 wild-type. Overnight cultures of the wild-type were inoculated into sterile potato tubers and extracted every 4 hours for a period of 24 hours (see materials and methods “Tissue maceration and total RNA extraction”). For statistical analysis of relative gene expression, the CT method was used (103). T-test was used to determine statistical significance (p-value <0.05).

### In planta competition assays in potato tubers

Competition assays were performed in potato tubers as described by Axelrood *et al.,* 1998 (104). In summary, *Solanum tuberosum* (cv. Mondial, a susceptible cultivar) tubers were sterilized with 10% sodium hypochlorite, rinsed twice with double distilled water, air-dried and then stabbed with a sterile pipette to 1cm depth. Overnight bacterial cultures of (*Pcb*1692 wild-type, *Pcb*1692Δ*pho*P, *Pcb*1692Δ*pho*P*-*p*pho*P and *Dickeya dadantii* with OD_300_ equivalent to 0.3) mixed in a 1:1 ratio and inoculated into surface sterilize potato tubers. Holes were sealed with petroleum jelly. Potato tubers were then placed in moist plastic containers and incubated for 24 hours at 25°C. Macerated tuber tissue was scooped out and the CFU/ml of surviving targeted bacteria determined by serial dilutions on LB supplemented with gentamycin (15µg/ml). The number of colonies observed was then converted to CFU/ml. This experiment was performed in triplicates, three independent times.

### Sequence analysis and protein domains characterization

Orthologous sequences of unannotated proteins (Big-associated and HrpV) from SRP strains were aligned using Clustal Omega (105). The resulting alignments were then used in iterative searches against the UniprotKB database (www.uniprot.org) through Hmmer search (32). The aligned results from the iterative searches were manually curated through Jalview alignment viewer (106), and the conserved areas in the alignments were selected. Next, those selected conserved blocks were analyzed for secondary structure prediction and programmatic gap removals by Jpred (107). The resulting concise alignments were submitted to HHpred (35) in order to identify appropriate templates for structure modelling. This strategy managed to retrieve several PDB structures to be used as templates. These templates were selected for subsequent modelling according to the predicted probability (>40%). For the Big-associated sequences, the following PDB entries were selected: 5K8G, 4N58, 4G75, 4FZL, 5K8G, 4FZM, and 2XMX; and 4G6T, 3KXY, 5Z38, 3EPU, 4GF3, 1JYA, 1S28, 5WEZ, 4JMF, and 1XKP selected for the HrpV. The models were predicted by Modeller (108), and visualization of these models generated through PyMOL (109).

### Horizontal gene transfer prediction and regulatory network analyses

Parametric methods are aimed to evaluate sequence composition bias in genes or genomic regions and measure their distance to the overall trend observed in the respective genome. These methods are especially powerful when applied in recent HGT candidates predictions contrarily to phylogenetic methods (110). Predictions were made with the support of two different methods, namely: GC content at the third codon position (GC3) and dinucleotide frequencies (DINT). The methods choice was adapted based on the conclusions drawn in the benchmark conducted by Becq, Churlaud (30). Genome-wide prediction of GC3 indexes of *Pcb*1692 and *Ddad*3937 coding sequences was made through the codonW tool http://codonw.sourceforge.net/). Dinucleotide frequency analyses were performed using the *fasta2kmercontent* script from the CGAT package (111). The Kullback-Leibler (KL) distance (112) was used to measure the difference between dinucleotide frequencies of individual coding genes and the respective genomes (averaged value of all coding sequences). KL distance is calculated according to the formula:

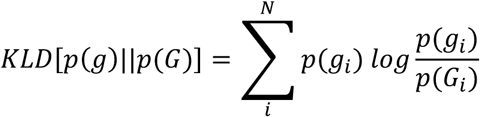

In which *p*(*g*) is the probability distribution of dinucleotide frequencies in an individual coding gene, and *p*(*G*) is the probability distribution of averaged dinucleotide frequencies from all coding genes in that same genome. KLD values were computed in R statistical environment. These two metrics (GC3 and DINT_KL) are then combined with the information on the genus-specificity of genes previously collected from orthologous clustering in order to predict HGT candidates. The above-threshold genes that were also genus-specific according to the previous analysis are predicted as HGT candidates. Gene networks were analyzed using Cytoscape (113). And the heatmap generated with Gitools (114).

## ACKNOWLEDGEMENTS

We thank Dr. Collins K. Tanui (former PhD candidate at the Department of Biochemistry Genetics and Microbiology) for the assistance in RNA extractions.

## SUPPORTING INFORMATION CAPTIONS

**S1.1 Table.** PhoP in planta regulon obtained from whole-transcriptome data set obtained at early infection (12 hours post-infection on potato tubers)

**S1.2 Table.** PhoP in planta regulon obtained from whole-transcriptome data set obtained at late infection (24 hours post-infection on potato tubers)

**S1.3 Table.** SlyA in planta regulon obtained from whole-transcriptome data set obtained at early infection (12 hours post-infection on potato tubers)

**S1.4 Table.** SlyA in planta regulon obtained from whole-transcriptome data set obtained at late infection (24 hours post-infection on potato tubers)

**S2 Table.** HGT prediction of genes of interest in *Pcb*1692 and *Ddad*3937 based on two parametric methods (dinucleotide frequencies and GC3 content)

**S3 Table.** Genome-wide detection of two newly discovered motifs in HrpV (GLLR) and Big-associated (GWYN) in 100 SRP genomes through HMMER scan

**S4 Table.** Gene-neighborhood of hrpV-containing genes in 89 SRP genomes represented by domain architectures predicted by HMMER3

**S1 Appendix.** Supporting material including S1-S5 Figures.

