## Supplementary material for "Horizontally acquired quorum sensing regulators recruited by the PhoP regulatory network expand host-adaptation repertoire in the phytopathogen *Pectobacterium carotovorum*": S1 Appendix

S1_Appendix - Supporting material including S1-S5 Figures.

SUPPORTING FIGURES

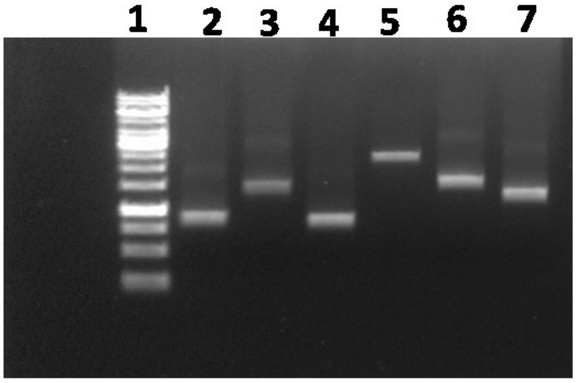


**Figure S 1 - PCR amplicons used to generate the phoP mutant.** Lane 1. DNA ladder, 2. *pho*P downstream PCR fragment, 3. kanamycin cassette PCR product, 4. *pho*P upstream PCR fragment, 5. Fusion product consisting of the downstream, kanamycin and upstream fragment, 5. Fusion product consisting of the downstream, kanamycin and upstream fragment. 6 & 7. Km1 and Km2 primers are internal kanamycin primers.

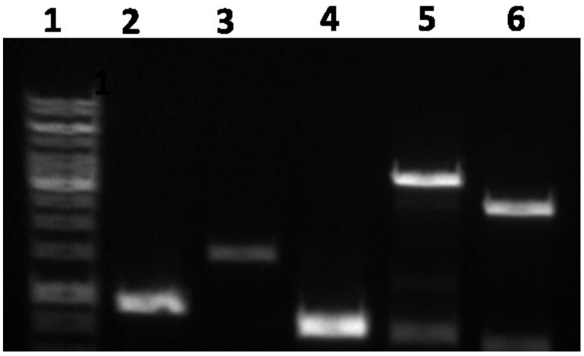


**Figure S 2 - PCR amplicons used to generate the *sly*A mutant.** Lane 1. DNA ladder, 2. *sly*A downstream PCR fragment, 3. kanamycin cassette PCR product, 4. *sly*A upstream PCR fragment, 5. Fusion product consisting of the downstream, kanamycin and upstream fragment. 6. Km1 primers, internal kanamycin primers.

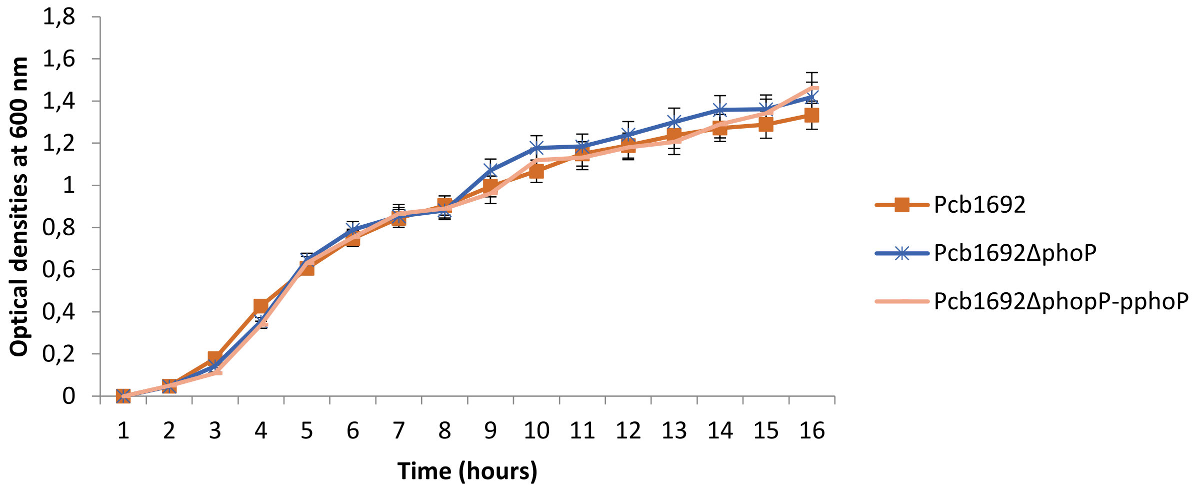


**Figure S 3 -** Growth and survival of *Pcb*1692 wild-type, *Pcb*1692Δ*pho*P and *Pcb*1692Δ*pho*P-p*pho*P in LB broth at 37°C for 16 hours with agitation at 370 rpm. Data points represent the means of three biological replicates.

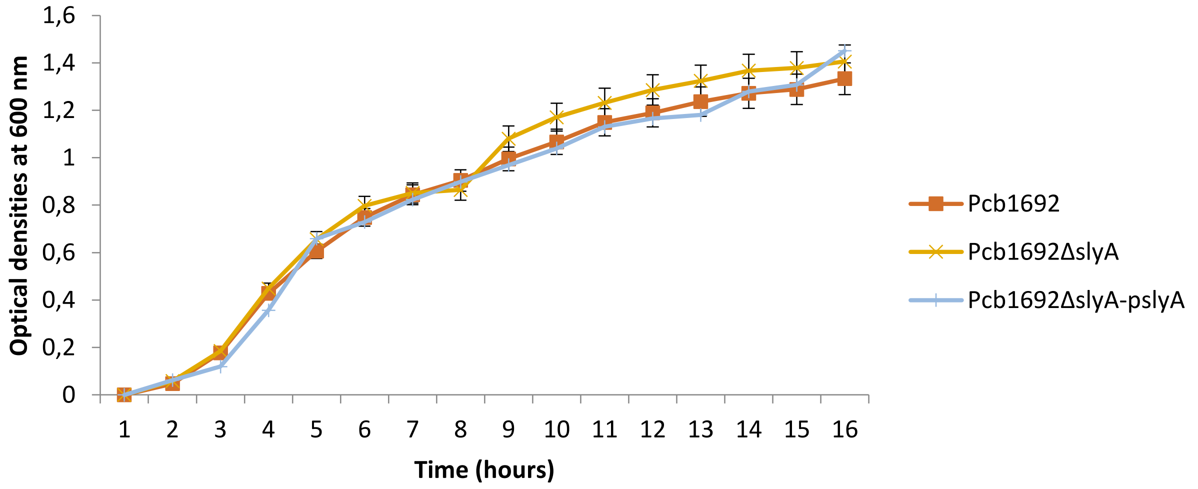


**Figure S 4 -** Growth and survival of *Pcb*1692 wild-type*,* *Pcb*1692Δ*sly*A and *Pcb*1692Δ*sly*A-p*sly*A in LB broth at 37°C for 16 hours with agitation at 370 rpm. Data points represent the means of three biological replicates.

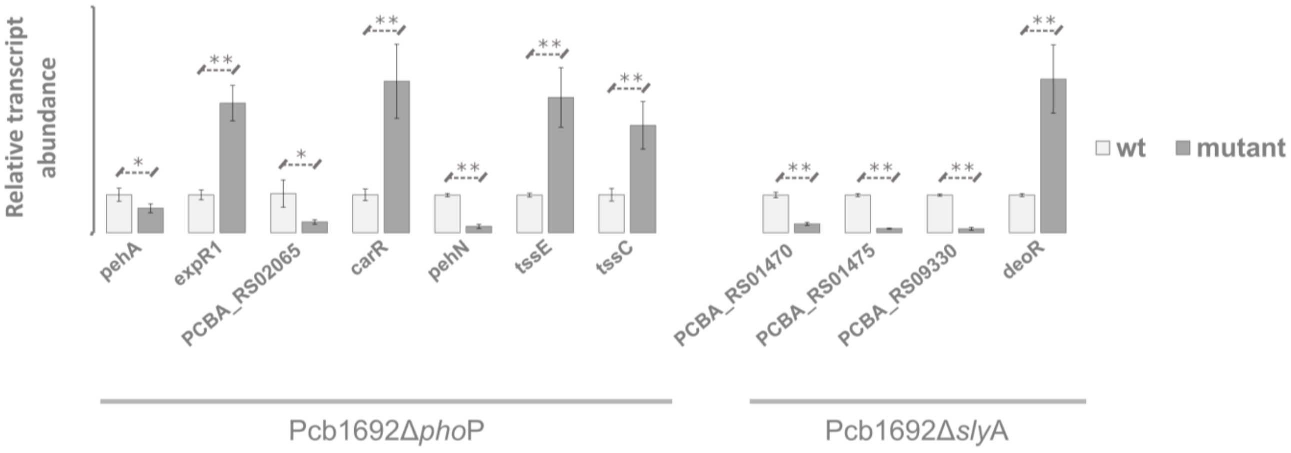


**Figure S 5 –** Additional validation of differentially expressed genes obtained from RNA-Seq. The relative transcript abundance was compared between mutant (Δ*pho*P or Δ*sly*A) and wild type (wt) strains of *Pcb*1692 through qRT-PCR. Statistical significance between wt and mutant strains for each gene was obtained through one-tailed T-test as follows: *($p > 0.05$); ** ($p > 0.01$). Genes annotated in the 'Carbohydrate metabolism' (KEGG: 09101) are represented by the respective NCBI locus tags. Genes associated with plant cell wall degradation (*peh*A – PCBA_RS10070; *peh*N – PCBA_RS15410), T6SS (*tss*E – PCBA_RS11165; *tss*C – PCBA_RS11175), and transcriptional regulation (*exp*R1 – PCBA_RS15665; *car*R – PCBA_RS04390; *deo*R – PCBA_RS02575) are represented by the gene names annotated through homology prediction using the eggNOG database.

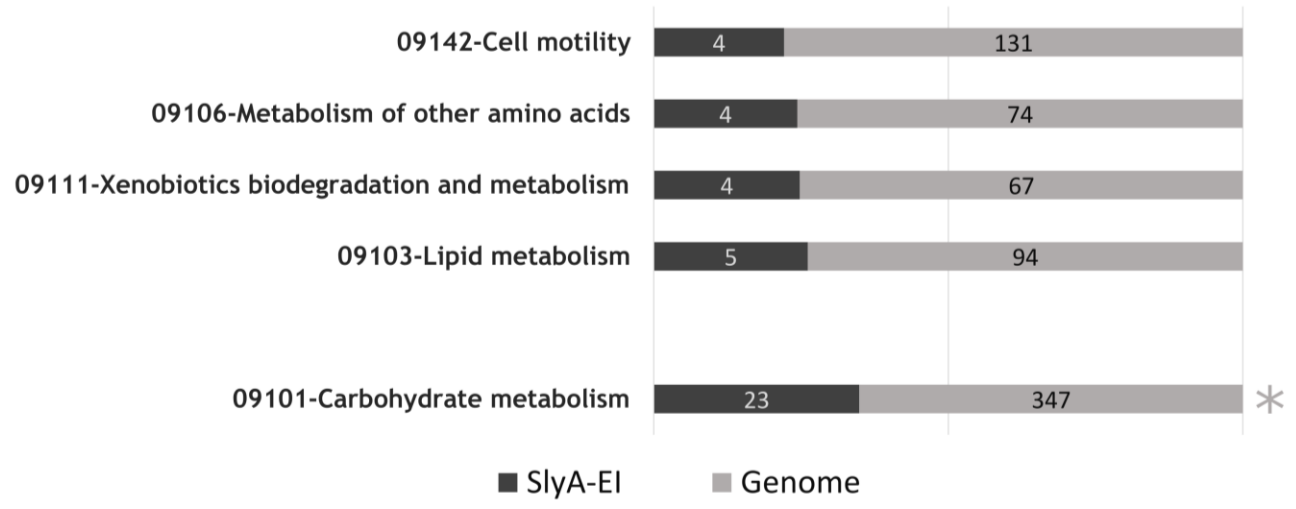


**Figure S 6 - Distribution of KEGG terms in the SlyA regulon at early infection.** The best represented KEGG terms within SlyA regulon at early infection are depicted in the graph according to their relative proportion (%) to the entire genome in a $\log_2$ normalized scale. A gap in the graph separates the categories found as enriched according to the adjusted p-value from Fisher exact tests at the bottom: *(FDR < 0.05).
